# A systematic analysis of gene-gene interaction in multiple sclerosis

**DOI:** 10.1101/2020.05.28.121939

**Authors:** Lotfi Slim, Clément Chatelain, Hélène de Foucauld, Chloé-Agathe Azencott

## Abstract

Multiple sclerosis is a complex autoimmune disease which genetic basis has been extensively investigated through genome wide association studies. So far, the conducted studies have detected a number of loci independently associated with the disease but few have investigated the interaction between distant loci, or epistasis. In the present work, we perform a gene level epistasis analysis of multiple sclerosis GWAS from the Wellcome Trust Case Control Consortium 2. We systematically study the epistatic interactions between all pairs of genes within 19 multiple sclerosis disease maps from the MetaCore pathway database. We report 4 gene pairs with epistasis involving missense variants, and 117 gene pairs with epistasis mediated by eQTLs. Our epistasis analysis is able to retrieve known interactions linked to multiple sclerosis: direct binding interaction between GLI-I and SUFU, involved in oligodendrocyte precursor cells differentiation, and regulation of IP10 transcription by NF-*κ*B, thus validating the potential of epistasis analysis to reveal biological interaction with relevance in a disease specific context.

## 1 Introduction

Extensive efforts have been deployed to tackle multiple sclerosis, a chronic disease damaging the central nervous system (Goldenberg 2012). A number of marketed drugs (Dargahi et al. 2017) attenuate the symptoms of the disease. However, an efficient drug targeting its root causes is still elusive. This is partially owed to our limited understanding of the mechanisms governing multiple sclerosis. Several studies demonstrated that heritability is a major component in multiple sclerosis (Dyment 2006; Dean et al. 2007). The development of GWAS has allowed to explore the genetic causes of this heritability. In GWAS, large cohorts of cases and controls are jointly studied in order to discover new biomarkers and causal loci. In the context of multiple sclerosis, at least fourteen studies(Sawcer, Franklin, et al. 2014) have been put in place in order to develop new hypotheses. So far, hundreds of loci (Baranzini and Oksenberg 2017; Cotsapas and Mitrovic 2018) have already been statistically associated with multiple sclerosis. The biology behind some of them (Gregory et al. 2007; Jager et al. 2009; Couturier et al. 2011) has been clarified while for the majority of retained loci, it remains unexplained (Sawcer, Franklin, et al. 2014).

GWAS in general, and in particular, the ones related to multiple sclerosis have enjoyed limited success (Dyment et al. 2004; Cotsapas and Mitrovic 2018) partially because of the used statistical methodology. Indeed, GWAS is classically conducted as a series of univariate statistical tests of association (Bush and Moore 2012) between a single-nucleotide polymorphism (SNP) and the phenotype. Though the statistical validity of this approach is indisputable, it suffers from a lack of statistical power because of high-dimensionality and multiple hypothesis testing (Shaffer 1995). It also suffers from a lack of interpretability due to the absence of a direct biological explanation for the significant SNPs. In addition, single-locus analyses, by design, do not take into account interactions between distinct genes, or epistasis (Phillips 2008). At least two gene-gene interactions have been discovered in multiple sclerosis: high levels of c-Jun may cause enhanced myelinating potential in Fbxw7 (Harty et al. 2019) and DDX39B is both a potent activator of IL7R exon6 splicing and a repressor of sIL7R (Galarza-Muñoz et al. 2017). An additional tripartite genic interaction has also been reported (Lincoln et al. 2009): epistasis between HLA-DRB1, HLA-DQA1, and HLA-DQB1 loci increases multiple sclerosis susceptibility. This further cements the need to study epistasis to understand the genetic basis of multiple sclerosis.

We perform here a selective gene-level analysis of epistasis in multiple sclerosis. The study of epistasis at the gene-level is important because the statistical association at the SNP level might not be strong enough to establish a link between the corresponding genes and the studied disease. We systematically study interactions between pairs of genes contained in 19 multiple sclerosis disease maps from the MetaCore (Ekins et al. 2006) dataset. For this purpose, we apply epiGWAS (Slim et al. 2018) on the multiple sclerosis GWAS from the Wellcome Trust Case Control Consortium 2 (Sawcer, Hellenthal, et al. 2011). EpiGWAS was originally developed for SNP-level detection, but we extended here to the gene-level. Our analysis yielded 4 gene pairs with epistasis involving missense variants, and 117 gene pairs with epistasis mediated by eQTLs. Among them, two pairs are already known: direct binding interaction between GLI-I and SUFU, involved in oligodendrocyte precursor cells differentiation, and regulation of IP10 transcription by NF-*κ*B. This confirms the capacity of the statistical study of epistasis to detect biological interactions that further our understanding of disease mechanisms.

## 2 Methods

### 2.1 epiGWAS: from the SNP level to the gene level

#### 2.1.1 Detecting SNP-SNP synergies with epiGWAS

In (Slim et al. 2018), we have developed epiGWAS, a new framework for targeted epistasis to detect interactions between a given SNP *A*, which we refer to as the target, and a set of SNPs *X* = {*X*_1_, · · ·, *X_p_*}, which can cover either the whole genome or a predetermined region e.g. a gene or a coding region. The output of epiGWAS is a set of interaction scores {*a*_1_, · · ·, *a_p_*} between each SNP in the set *X* = {*X*_1_, · · ·, *X_p_*} and the target *A*. We propose a family of methods to compute the interaction scores. All interaction scores account for the relationship between the target and the rest of the genome through a propensity score *π*(*A*|*X*). This propensity score models the linkage disequilibrium structure between *A* and *X*. Taking it into account allows us to account for main effects and recover epistatic effects only.

If we choose a symmetric binary encoding for the target *A* ∈ {−1, +1}, we can always write the following decomposition for the genotype-phenotype relationship:

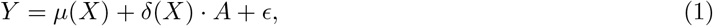

where *ϵ* is a zero mean random variable and

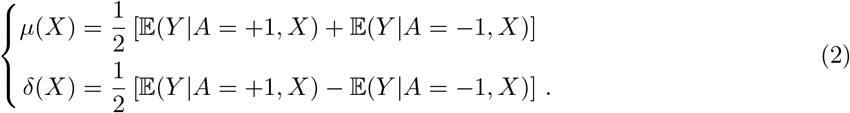

The first term in Eq. 2, *μ*(*X*), models the average effect of the target *A* on the expected phenotype, conditionally on *X*. By contrast, the second term *δ*(*X*) models the difference of the expected outcome for the two modes of *A* ∈ {−1, +1}. The term *δ*(*X*) explicits any conditional effect of *A* that can not be solely explained through the SNPs in *X*. This why we interpret the product term *δ*(*X*) · *A* as an interaction term between *A* and *X*.

For a given sample, only one of the two possibilities {−1, +1} is observed. This makes directly estimating the term *δ*(*X*) impossible. The purpose of epiGWAS is to introduce propensity scores to recover the term *δ*(*X*). More precisely, we are interested in recovering the support of *δ*(*X*), namely the SNPs within *X* interacting with *A*.

We notice that by using a second binarized version of the target, 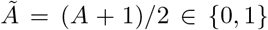, we can directly derive the desired term *δ*(*X*):

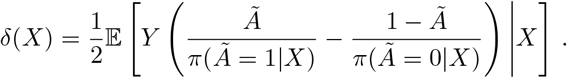

So, a first straightforward approach is to implement a penalized regression approach for the estimate of the support of *δ*(*X*). We refer to this approach as *modified outcome*. We use this denomination because the natural outcome *Y* is substituted by the modified outcome:

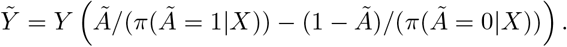

To derive 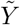, an estimate of the propensity score *π*(*A*|*X*) is needed. The classical and straightforward approach for the estimation of *π*(*A*|*X*) is logistic regression. For genomic data, given the high dimensionality of *X*, we observed that extreme overfitting ensued. As a solution, we resorted instead to a semi-parameterized estimation method. In fastPHASE (Scheet and Stephens 2006), a hidden Markov model (HMM) is developed in order to perform imputation. The observed states correspond to SNPs and the hidden states to structural dependence states. After fitting the HMM model in a chromosome-wise fashion, we applied the forward algorithm (Rabiner 1989) to obtain the scores *π*(*A*|*X*).

If the estimation error of *π*(*A*|*X*) is large or severe overfitting occurs, the use of the inverse of the estimated scores can result in numerical instability and bias the results. Several approaches have already been proposed in the literature (Lunceford and Davidian 2004) to tackle this issue.

Among them, we only use the *robust modified outcome* method. In a previous work (Slim et al. 2018), we have demonstrated its superior performance in comparison with other epistasis detection baselines and the other methods of the modified outcome family.

#### 2.1.2 Gene-level epiGWAS

EpiGWAS can be ran in an exhaustive fashion for each target *X_i_* against the rest of the SNPs {*X*_1_, · · ·, *X_i−_*_1_, *X_i_*_+1_, · · ·, *X_p_*}. This procedure generates a list of interaction score vectors. The interpretability and usability of such an output is limited because of the large number of interactions and the different covariates for each target which makes the comparison of the associated scores difficult. For instance, different regularization grids yield different stability curves, and thus, different areas under the curve. Furthermore, despite their robustness, the biological significance of the scores is limited. A first step to improve interpretability is to use rankings. From a practical point of view, rankings are a sensible choice because only the highest-ranking SNPs are used. Rankings also improve comparability between different targets because of the similarity of scale and insensitivity to the underlying parameterization. For a target *i*, we denote *r_ij_* ∈ {1, · · ·, *p* − 1} the rank in a decreasing order of the score of SNP *j*.

Another immediate benefit of the use of rankings is the possibility of combining of different rankings. For example, for two SNPs *i* and *j*, we can define the following epistasis interaction score:

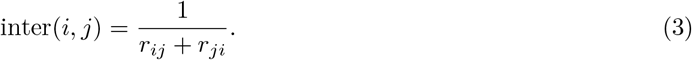

The interaction score in Eq. 3 has the advantages of symmetry and boundedness. The scores are comprised between [0, 1/2]. Additionally, the combination of two pairwise scores *r_ij_* and *r_ji_* can help control the estimation errors for one of the targets. For example, if two SNPs *i* and *j* are in interaction and the result *r_ij_* is not sufficiently high to reflect that, a good ranking of *r_ji_* can help compensate that.

We can further aggregate the rankings to detect interactions between genes. More generally, the rankings can be combined to detect interactions between any disjoint sets of SNPs *e.g.* biological pathways, regulatory regions, etc. Let *p′* be the total number of genes and {*G*_1_, · · ·, *G_p′_*} the corresponding sets of SNPs such that 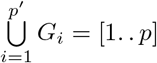. The easiest way to devise an interaction score between two genes *i′* and *j′* is to compute the average of all pairwise scores:

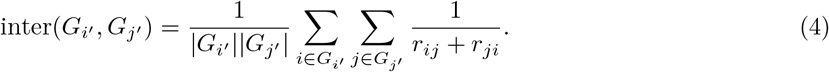

Thanks to the symmetry of SNP-SNP scores in Eq. 3, the gene-gene scores in Eq. 4 are symmetric, too. Moreover, the averaging reduces the impact of the size of the genes. In addition to the mean, we can also use the median or the minimum/maximum of all pairwise scores. However, only a single value will be taken into account with the latter strategies. Depending on the implemented regression method, with respect to a target *i*, the scores, and hence the rankings, of two nearby variants *j* and *j′* can be similar because of linkage disequilibrium. This can make the gene-gene scores more robust through the averaging of high nearby rankings. On the other hand, the averaging strategy can be biased by the marginal effects of some *loci* inflating by consequence the interaction scores.

### 2.2 Data and experiments

In this section, we describe the data we integrate to perform our systematic gene-gene interaction analysis for MS. For genotypic data, we select the MS dataset from the second release of the Wellcome Trust Case Control Consortium (WTCCC2) et al. (Sawcer, Hellenthal, et al. 2011). In order to improve statistical power and the downstream biological interpretation, we subset the marker SNPs related to the genes referenced in the MetaCore (Ekins et al. 2006) disease maps for multiple sclerosis. Each gene pair within a disease map is tested for interaction. Within the same disease map, the included genes affect the same MS-related mechanism. Therefore we can use this prior knowledge to evaluate if our method can retrieve known interactions and identify new ones. The SNPs can be mapped to the genes in two different ways:

- Physical mapping: we select all the marker SNPs which positions are within the boundaries of a gene. In this case, we take into account SNPs with an effect on the structure and function of the corresponding protein.
- eQTL-SNP mapping: with the selection of eQTL SNPs, we study epistasis through the variation in expression of the associated genes.

### 2.3 Genotypic data

The WTCCC2 study includes 9 772 MS cases and 17 376 controls hailing from 15 different countries. The presence of population structure, confirmed by a genomic inflation factor (GIF) of 3.72, is poised to lead to inference issues. To avoid this problem, we only use Caucasian British samples in both cases and controls. The resulting dataset consists of 2048 cases and 5733 controls with a GIF of 1.06 which proves the homogeneity of the dataset. The selected controls come from two distinct cohorts from the UK Blood Services (NBS) and the 1958 British Birth Cohort (58C). The careful reader may notice the important imbalance between the total number of cases and controls which may distort the results. To equalize the field, we randomly subsample controls to obtain a number of controls equal to the number of cases. We also note that we discarded the samples singled out for quality control by the WTCCC.

#### 2.3.1 Variant selection

We give in Table 1 the full list of MS disease maps. For ease of reproducibility, we also give the internal ID of the disease maps, as indicated in MetaCore. The number of genes within each map greatly varies. It ranges from 13 genes for disease map (DM3305) to 100 genes (DM4593). Even for the larger maps, the total number of genes is still low enough to perform exhaustive pairwise analysis for all SNPs mapped to the selected genes. Similarly for sample-wise QC, we first discarded all low quality SNPs designated by the WTCCC2. We then selected SNPs according to the following mappings:

**Table 1.** Titles and internal IDs of MetaCore disease maps related to MS.

| internal ID | Title |
| --- | --- |
| 3302 | Notch signaling in oligodendrocyte precursor cell differentiation in multiple sclerosis |
| 3305 | SHH signaling in oligodendrocyte precursor cells differentiation in multiple sclerosis |
| 3306 | Inhibition of oligodendrocyte precursor cells differentiation by Wnt signaling in multiple sclerosis |
| 4455 | Inhibition of remyelination in multiple sclerosis: regulation of cytoskeleton proteins |
| 4593 | Axonal degeneration in multiple sclerosis |
| 4693 | Role of Thyroid hormone in regulation of oligodendrocyte differentiation in multiple sclerosis |
| 4703 | Demyelination in multiple sclerosis |
| 4791 | Role of CNTF and LIF in regulation of oligodendrocyte development in multiple sclerosis |
| 4794 | Retinoic acid regulation of oligodendrocyte differentiation in multiple sclerosis |
| 4843 | Growth factors in regulation of oligodendrocyte precursor cells proliferation in multiple sclerosis |
| 4846 | Growth factors in regulation of oligodendrocyte precursor cells survival in multiple sclerosis |
| 4901 | Inhibition of remyelination in multiple sclerosis: role of cell-cell and ECM-cell interactions |
| 5199 | Cooperative action of IFN-γ and TNF-α on astrocytes in multiple sclerosis |
| 5288 | Impaired inhibition of Th17 cell differentiation by IFN-β in multiple sclerosis |
| 5378 | Role of IFN-β in the improvement of blood-brain barrier integrity in multiple sclerosis |
| 5398 | Role of IFN-β in activation of T cell apoptosis in multiple sclerosis |
| 5518 | Role of IFN-β in inhibition of Th1 cell differentiation in multiple sclerosis |
| 5601 | IL-2 as a growth factor for T cells in multiple sclerosis |
| 5611 | Role of IL-2 in the enhancement of NK cell cytotoxicity in multiple sclerosis |

- Physical mapping: corresponds to retrieving all marker SNPs located on a given gene. We use the accompanying R package metabaser (Ishkin 2019) to first define the boundaries of a given gene, and then subset all SNPs according to their positions, as referenced in dbSNP version 144 (Pagès 2017).
- eQTL mapping: we use the cis-eQTL dataset from the eQTLGen consortium (Võsa et al. 2018), which provides for each gene a list of significant eQTL-SNPs. The dataset combines 31 684 whole blood samples from 37 cohorts.

For our present study, we chose cis-eQTLs instead of trans-eQTLs because of their higher degree of association to gene expression. The higher association can be attributed to the proximity of the SNPs to the genes: cis-eQTL are located within 1 Mb window from a gene and they often closely map to either the transcription start site or the transcription end site of a gene. The application of a false discovery rate (FDR) of 0.05 resulted in the identification of eQTL-SNPs for 16 989 genes, or approximately 88.3% of all autosomal genes expressed in blood and tested in the cis-eQTL analysis. We restricted ourselves to the genes present in the metaCore disease maps. We observed that the obtained eQTL-mapping datasets were larger than the physical mapping datasets in terms of number of SNPs: the median number of SNPs per disease map is 392 for the physical mapping analysis and 999 for the eQTL-mapping analysis. In Appendix A, we give the exact number of SNPs per disease map for each type of mapping. We also included the average number of SNPs per gene for each disease map and for both mappings.

Even though the two analyses are unrelated and use different sets of SNPs, some concordance for the top-scoring genes is to be expected. In fact, for the eQTLGen consortium, (Võsa et al. 2018) show that out of 15 317 trait-associated SNPs, 15.2% were in high LD with the lead eQTL SNP showing the strongest association for a cis-eQTL gene. Although the mentioned association is far from perfect, it demonstrates the often-overlooked link between the two analyses.

## 3 Results

We exhaustively apply our gene-gene interaction scores in Eq. 4 to obtain *p′*(*p′* − 1)/2 interaction scores per disease map, where *p′* is the number of genes. Given the size of the maps (see Appendix A), the interpretation of the full results is rather difficult. We instead focused on the 2% top-scoring pairs for the two analyses. The 2% threshold was manually set with respect to the obtained result. We remarked that the top-scoring edges often constituted connected sub-components. We also remarked that the obtained sub-components for the eQTL and physical mappings are often interlinked. These two remarks are more commented in the following paragraphs. We give an illustration of the results in Figure 1 in which we plot the obtained subnetworks in addition to the original edges for DM 3306. We relegate the results of the other disease maps to Appendix B.

**Figure 1.**
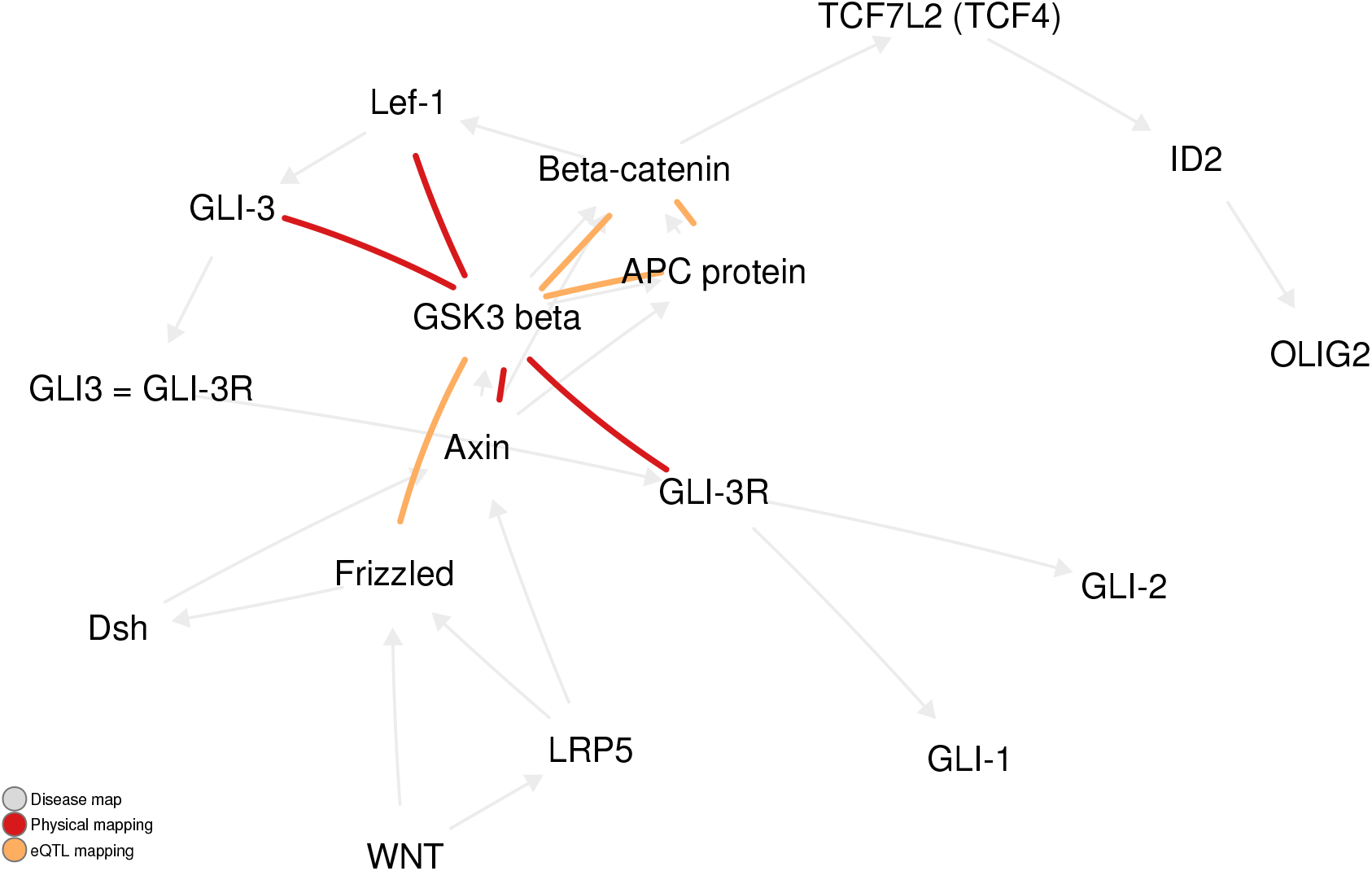
The 2% top-scoring pairs in DM 3306 for eQTL and physical mappings.

We notice a general consistency of the results between the different disease maps, which can be formulated through the characteristics below. We also conduct an enrichment analysis, from which we derive empirical *p*-values to measure the statistical significance of the observed characteristics (see Appendix C for the full results).

- Connectedness: the obtention of connected components for both mappings is the most important aspect of the results. With the exception of DM 3305, 3306 and 4794 consisting of 1 or 2 edges, all disease maps have a *p*-value lower than 0.05. Of particular interest are large components because of their significance. In many cases, we obtained an empirical *p*-value of 0 despite using 10^4^ simulations. The discovery of these novel subnetworks can help the understanding of multiple sclerosis by unraveling new disease mechanisms.
- Complementarity: with the exception of disease map 4593, the subnetworks of the two mappings are connected *i.e.* they share at least one common node. In fact, they are often connected through multiple nodes without a significant overlap between the edges of the two networks. For instance, they share 5 vertices in DM 4901. In Appendix C, we quantify the significance of having 1, 2 or 3 genes in common. We particularly note that 3 edges are in common in DM 3302 for a *p*-value of 0.038. Therefore, the two types of mappings recover distinct, though connected, interactions, which suggests the complementarity of the two mappings. We can then consider the union of the two subnetworks for further study.
- Centrality: we observed a high degree of connectivity for certain nodes. For example, we mention FAK in DM 4901 (*p*_FAK_ = 0), SHP-2 in DM 4843 (*p*_SHP-2_ = 0.014) and TRADD in DM 4843 (*p*_TRADD_ = 0.052). We attribute this centrality to the existence of important marginal effects that were not completely filtered out. Interestingly, the role of these genes in MS has already been established (Sun et al. 2010; Ahrendsen et al. 2017; Reuss et al. 2014).
- Commonality: despite using the top 2% of all *p′*(*p′* − 1)/2 possible edges for each disease map, some of the retained edges were already present in the original disease maps. In at least 9 out of 19 disease maps, a single edge already exists in the original disease map, and in at least four of them two edges. In DM 3306, we even recover three edges (*p* = 0.099). Nonetheless, drawing conclusions about the underlying biology is challenging given the potential mismatch between biological epistasis and statistical epistasis (Moore and SM Williams 2005).

### 3.1 Enrichment analysis for obtained subnetworks

Beyond the validation with existing edges, the main goal of the systematic analysis we conduct here is to discover novel gene-gene interactions in multiple sclerosis. Their biological validation requires laboratory experiments to confirm the observed statistical synergy. As we do not have access to such facilities, we use the enrichment of the recovered networks in terms of existing therapeutic targets as a validation metric. The chosen metric can be criticized in two ways: it is biased in the sense that therapeutic targets only reflect our current understanding of the disease and the existence of effective molecules for the targets. In addition, the targets were often selected on an univariate basis, while the subject of the current study are epistatic interactions. However, an enrichment analysis in terms of therapeutic targets has the advantages of being a trustworthy background thanks to the proven effect of the included genes and its relevance in terms of development of future therapies. For instance, combination therapies if an existing therapeutic target is shown to be interacting with another gene within the recovered subnetworks. Moreover, in light of the new FDA guidance for the co-development of two or more drugs ^I^, our study pipeline can be of special interest because of its focus on synergistic effects instead of separate additive effects.

In our case, we use OpenTargets (Carvalho-Silva et al. 2018a) as a dataset for therapeutic targets. The dataset is a collaborative effort to create an up-to-date and comprehensive repository to link genomic information of drug targets to a disease of interest. The enrichment analysis studies the overpresence of OpenTargets targets in the obtained networks in comparison with the original disease maps. We use for this matter a classical hypergeometric test (Rivals et al. 2006) to determine the statistical significance of their overpresence. We give the resulting *p*-values in Appendix D. For twelve disease maps, we found at least one common gene between our subnetworks and OpenTargets. Given a significance threshold of 0.05, we found two significant disease maps DM 4593 and DM 5378 with respective *p*-values of 0.008 and 0.02. The enriched subnetworks require further investigation, especially to study the links within the known targets and between the known targets and the rest of the subnetwork.

### 3.2 Directionality of the synergy

As shown before, our gene-level pipeline with epiGWAS robustly detects the presence of epistatic synergies between two genes. However, the obtained interaction scores do not allow to determine the directionality of the synergy. The synergy can be either positive or negative by respectively increasing or decreasing the disease risk probability. We can nonetheless get a partial answer by studying the nature of interaction between the top-scoring SNPs for each gene pair. We only selected the top-scoring pair because of its disproportionate impact on the corresponding gene-gene score. For example, we can consider the extreme case where for a pair of SNPs (*i, j*), we have *r_ij_* = *r_ji_* = 1. The next possible best scoring pair is *r_i′j′_* = *r_j′i′_* = 2 and it further decreases in a hyperbolic manner for the lower rank pairs. So, in the best cases, the top pair will be at least twice as important as the following one.

The direction of the synergy between two uni-dimensional variables can be studied in various ways (VanderWeele and Knol 2014). In particular, for a binary outcome *Y* and two variables *X*_1_ and *X*_2_, we can study the sign of the interaction coefficient *α*_12_ in the following logistic model: logit *P* (*Y* |*X*_1_, *X*_2_) = *α*_0_ + *α*_1_*X*_1_ + *α*_2_*X*_2_ + *α*_12_*X*_1_*X*_2_. Logistic models are widely used for the study of epistasis. For the physical mapping strategy, we conduct a similar analysis. As for the eQTL mapping strategy, the methodology we use for physical mapping can be refined to amount to the desired gene-level interactions. The effect of a SNP *i* on the expression level *e_i_* of the corresponding gene *G_i_* can be examined through a model of the form *e_i_* = *γ_i_* + *β_i_X_i_*. The directionality of the synergy can be deduced from the sign of the following ratio:

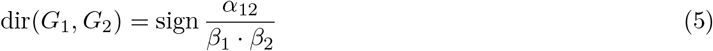

To get a better grasp of the meaning of the score in Eq. 5, it suffices to replace the two linear expression models directly in the interaction logistic model. Precisely, we obtain:

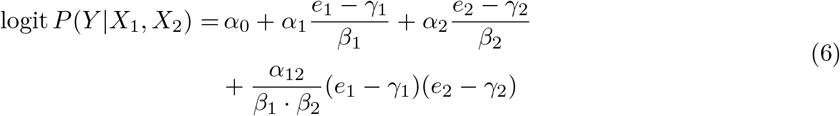

The synergy of the two gene expressions is given by the coefficient *α*_12_/(*β*_1_ · *β*_2_) which sign determines the directionality of the epistatic interactions between the two genes. To the best of our knowledge, this is the first study which studies epistasis from such a perspective by including eQTL scores in this way and by moving back and forth between SNP-level and gene-level epistasis. Furthermore, the synergy score in Eq. 5 can also be interpreted as an extension of Mendelian randomization (Davies et al. 2018) to second-order interaction effects.

The eQTLGen consortium (Võsa et al. 2018) does not directly supply the effect sizes *β*_1_ and *β*_2_ in the linear expression models. For each SNP, the effect size *β* is derived from the corresponding Z-score using the following relationship:

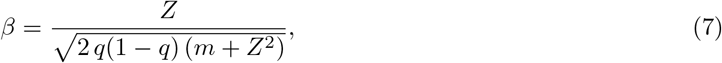

where *q* is the MAF of the SNP of interest, as reported in the 1kG v1p3 ALL reference panel and *m* is the cohort size.

For the significant interactions, we provide a csv file containing the list of coefficients *α*_12_ in addition to (*m*_1_, *q*_1_, *Z*_1_), (*m*_2_, *q*_2_, *Z*_2_) and the directionality of the synergy dir(*G*_1_, *G*_2_) ∈ {−1, +1} for the eQTL strategy. One possible approach to appraise the results is to consider a number of summary statistics to get an overview of the kind of synergies occurring within biological pathways. Interestingly, for all SNP pairs, the interaction coefficient *α*_12_ is positive in 47% of all cases and the directionality of the synergy dir(*G*_1_, *G*_2_) is equally split between positive and negative. For the eQTL strategy, we found that *α*_12_ and dir(*G*_1_, *G*_2_) agree approximately half of the time (48%). This gives further credence to our gene-gene approach by showing that a different type of information can be obtained by considering more biologically-relevant gene-level interactions.

For each SNP, we also include its PolyPhen (Adzhubei et al. 2013) and SIFT (Ng 2003) scores reported in BioMart (Kinsella et al. 2011) to better understand its potential deleterious impact on MS. If available, both scores are comprised between 0 and 1, but with opposite interpretations. For SIFT, 0 denotes a deleterious amino-acid substitution, while for PolyPhen, 1 denotes an benign substitution. In total, we obtained 5 variants which were predicted as deleterious by at least one of the two methods.

### 3.3 Biological interpretation

In addition to the preceding statistical analysis, we also conduct a biological analysis of the results for both mappings. Our analysis is built upon existing information in MetaCore disease maps in conjunction with relevant literature.

#### 3.3.1 Physical mapping

In total, we obtained 136 epistatic interactions in the 19 disease maps. As an exhaustive analysis of all interactions is out of reach, an a posteriori filtering is needed. In physical mapping, an epistatic interaction between two genes corresponds to a change of their protein structure. We therefore retain an interaction if at least one of the SNPs in the top-scoring pair can lead to a loss of function at the protein level. For that matter, the SNPs are selected according to the following criteria:

- Frameshift variant or incomplete terminal codon variant or missense variant or start loss variant,
- Stop-gained, stop-lost or stop-retained variant,
- Terminal codon variant.

The filtering process yielded 4 gene pairs where one of the the genes presents a missense variant (Appendix G). For each of these gene pairs, the impact on the MS phenotype is given as specified (activation or inhibition) or unspecified (unknown), as depicted in Fig 2. Among the obtained 4 pairs, GLI-1 and SUFU appear to be particularly interesting, since both genes are in direct binding interaction in DM 3305, which illustrates the SHH (Sonic Hedgehog) signaling in oligodendrocyte precursor cells differentiation in MS (Appendix E.1).

**Figure 2.**
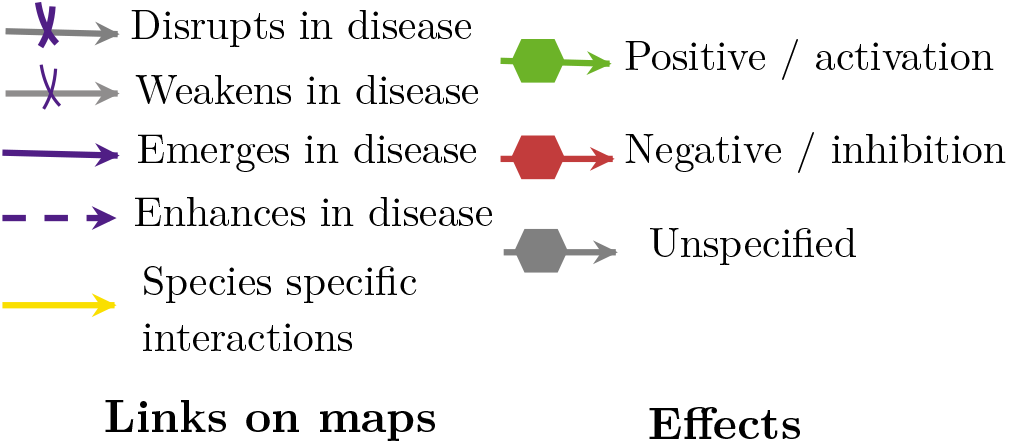
The different types of links between proteins/proteins or proteins-phenotypes in MetaCore maps

#### 3.3.2 eQTL mapping

In eQTL mapping, an epistatic interaction consists of a gene pair, the simultaneous up/down-regulation of which induces a synergistic effect which lowers or increases the risk of MS. To better understand the impact of simultaneous gene up-regulation on disease propensity, we differently rewrite Equation 6:

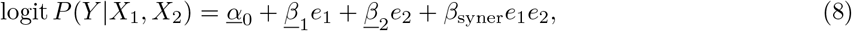

where *β*_syner_ = *α*_12_/(*β*_1_ · *β*_2_) and the constants *α*_0_, *β*_1_ and *β* _2_ are functions of (*α*_0_, *α*_1_, *α*_2_, *α*_12_), (*γ*_1_, *γ*_2_) and (*β*_1_, *β*_2_).

The impact of gene up-regulation can be assessed through the signs of (*β*_1_, *β*_2_, *β*_syner_). For instance, if *β*_1_, *β*_2_ and *β*_syner_ are positive, an increase in the expression of either genes leads to a higher disease risk. Hence, a joint inhibition of the two genes reduces the risk. In Table 2, we similarly study all possible sign combinations of (*β*_1_, *β*_2_, *β*_syner_) to devise a number of recommendations for the application of epistasis to the development of combination therapy.

**Table 2.** Analysis of the impact of genes up-regulation on the risk for humans to develop MS, for each gene individually (signs of *β*_1_ and *β*_2_), and for the pair of genes synergistically (sign of *β*_syner_) which is epistasis.

| $\beta_1$ | $\beta_2$ | $\beta_{\text{syner}}$ | Impact of $\beta_1$ and $\beta_2$ on MS | Recommendation for combination therapy |
| --- | --- | --- | --- | --- |
| $> 0$ | $> 0$ | $> 0$ | detrimental | inhibition of the two genes reduces the risk for MS |
| $> 0$ | $> 0$ | $< 0$ | beneficial | genes must not be inhibited |
| $< 0$ | $< 0$ | $< 0$ | beneficial | genes could be activated at the same time |
| $< 0$ | $< 0$ | $> 0$ | detrimental | genes must not be activated |
| $> 0$ | $< 0$ | NC | NC | NC |

A total of 117 gene pairs in 19 disease maps were obtained with the eQTL mapping strategy. As in physical mapping, an additional filtering is needed. We selected the gene pairs in which the coefficients (*β*_1_, *β*_2_, *β*_syner_) share the same sign (all positive or negative). If positive, the inhibition of both genes reduces the risk for MS. By contrast, if negative, the two genes should be jointly activated to reduce MS risk. This filtering led to 25 gene pairs of interest across 13 maps. Since a thorough study of all 25 pairs is impossible, we implemented an additional filtering criterion: existence of a specified effect on MS-related phenotypes e.g. demyelination, remyelination failure, oligodendrocyte death, damage of neural axons, etc. The effect nature is given by the arrow types (see Figure 2). This final filter led to 9 gene pairs to consider (see Appendix F).

Confident in the single gene pair where both genes have a specified impact on the phenotype, NF-*κ*B and IP10 (see Appendix H), we have investigated in further details their role in MS in the aim of assessing their synergistic effect on MS physio-pathology. Our analysis is focused on DM 5199 (see Appendix E.3) where both genes belong to essential pathways.

##### Role of IP10 in MS: recruitment of T cell in the CNS

IP10 (or IP-10 / CXCL10 (C-X-C motif chemokine ligand 10) / Interferon-Inducible Cytokine IP-10) is an antimicrobial gene which encodes a chemokine of the CXC subfamily, and is a ligand for the receptor CXCR3. This pro-inflammatory cytokine is involved in a wide variety of processes such as chemotaxis, differentiation, and activation of peripheral immune cells, like monocytes, natural killer, T-cell migration, and modulation of adhesion molecule expression (Romagnani et al. 2001; Antonia et al. 2019; Tokunaga et al. 2018).

IP-10 is strongly induced by IFN-*γ* as well as by IFN-*α*/*β* (Qian et al. 2006). In vitro, CXCL10 can also be induced by NF-*κ*B, and has been shown to have an early role in hypoxia-induced inflammation (Schmid et al. 2006; Xia et al. 2016). Indeed, in the disease map, the activation of IP10 by NF-*κ*B is clearly indicated by an activation arrow (green arrow). Thus, the two genes are in direct interaction, where NF-*κ*B regulates the transcription of IP10.

DM 5199, which contains IP10 and NF-*κ*B, is focused on the impact of beta-2 adrenergic receptors, which are lacking in astrocytes in MS. This lack enables IFN-*γ* and TNF-*α* to trigger the expression of several key pro-inflammatory genes (Keyser, Zeinstra, et al. 2004; Keyser, Laureys, et al. 2010). Whereas human astrocytes are only partially competent antigen presenting cells, the upregulation of MHC-II by IFN-*γ* alone or in combination with TNF-*α* enables astrocytes to present myelin as an auto-antigen, and triggers the production of the co-stimulatory molecules C80 and CD86 at their surface. Experimentally, the expression of MHC-class I and MHC-class II, together with the co-stimulatory molecules CD80 and CD86, is detectable in astrocytes in MS plaques (TRAUGOTT and LEBON 1988).

After the transformation of astrocytes in immuno-competent cells, IP10 plays a major role by activating the recruitment of Th1 cells into the CNS (Fig 3a). Indeed, in MS, activated CXCR3+ T-cells (IP10 is the ligand for the receptor CXCR3) enter the CNS, and can be located in the cerebrospinal fluid or in the brain parenchyma (Lassmann and Ransohoff 2004). This transport is made possible due to the blood Brain Barrier disruption in MS (Minagar and Alexander 2003).

**Figure 3.**
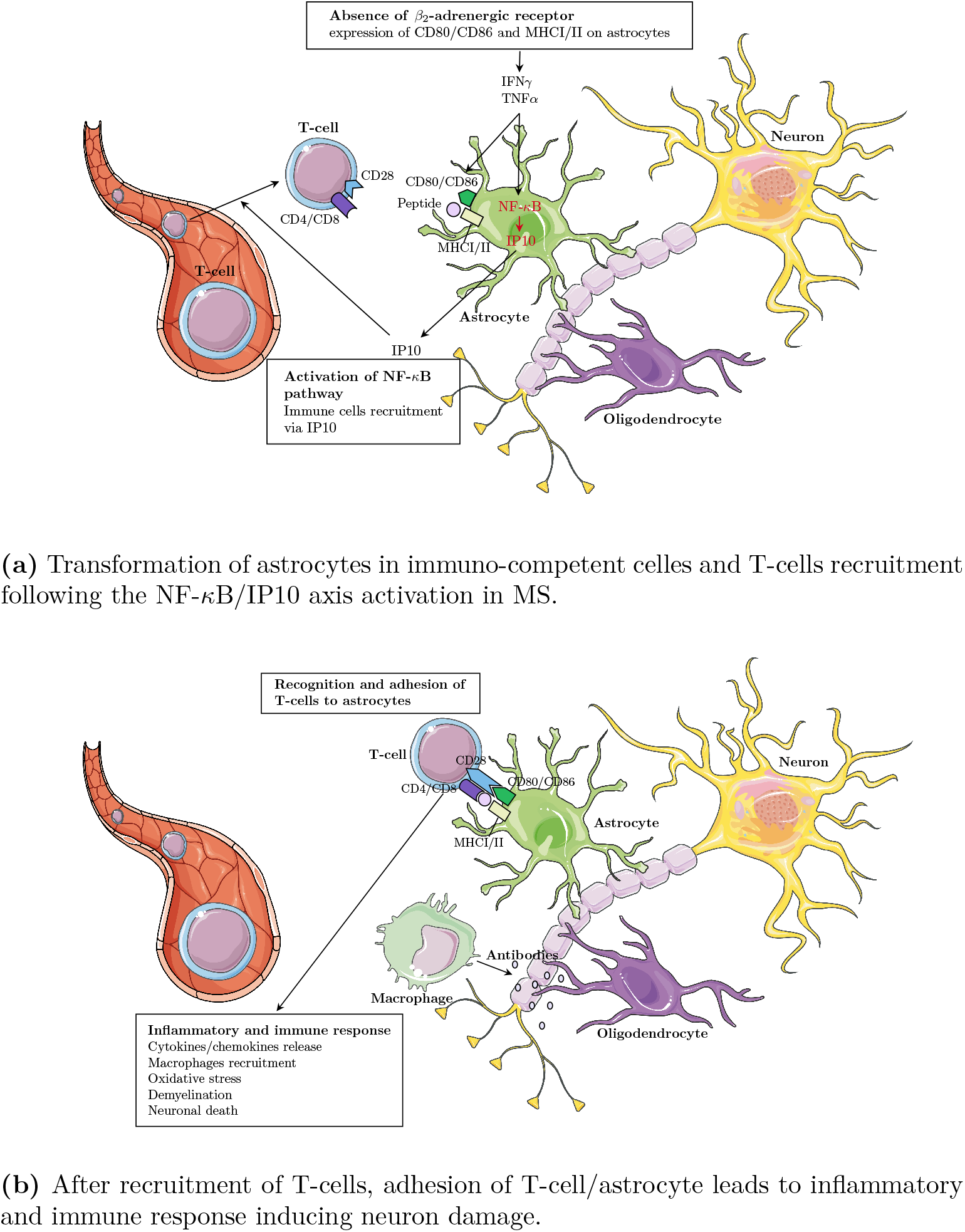
Schematic representation of the role played by the gene pairs NF-*κ*B/IP10 in the development of demyelination in MS.

Arriving in the CNS, T lymphocytes recognize astrocytes via their MHC-II, and anchor them via their CD28 which binds to CD80 and CD86 on astrocytes. This intercellular contact between T cells and astrocytes presenting myelin antigens induces the reactivation of T cells in the CNS (Cornet et al. 2000). T cells then secrete pro‐inflammatory cytokines; demyelination occurs and macrophages are activated. This further damages myelin and releases cytokines - but also phagocytosing myelin debris - which leads to the damage of neural axons (A Williams et al. 2007) (see Fig 3b).

##### Role of NF-*κ*B in MS: transcription regulation

Astrocyte reactivity is regulated by key canonical signaling cascades, among which the NF-*κ*B pathway is qualified as pivotal for establishing neuroinflammation (Ponath et al. 2018). TNF-*α* binds to TNF-R1, which is constitutively expressed in astrocytes, and activates NF-*κ*B signaling pathway (Liang et al. 2004). In cytoplasm, NF-*κ*B is inhibited by I-kB proteins. Phosphorylation of I-*κ*B by IKK (cat) kinase complex marks I-kB for destruction via the ubiquitination pathway, thereby allowing activation of NF-*κ*B complex (Liang et al. 2004). The activated NF-*κ*B translocates into the nucleus and upregulates transcription of target genes including IP10 (Majumder et al. 1998).

##### Status of IP10 and NF-*κ*B as potential targets in MS treatment assays

Human IP10 is a secreted protein, and is mainly located in the extracellular space, but also in the plasma membrane, and to a lesser extent in the cytosol and nucleus (Source: UniProtKB/Swiss-Prot). Today, the ChEMBL database indicates that two antibodies of IP10 are studied in clinical trials: NI-0801 (Phase I completed for allergic contact dermatitis, Phase II terminated for primary biliary cirrhosis) and ELDELUMAB (phase II mainly for rheumatoid arthritis, ulcerative colitis and Crohn’s disease; source: Open Targets (Carvalho-Silva et al. 2018b)). The fact that, except for allergic contact dermatitis, all of these diseases belong to the auto-immune diseases family like MS, suggests that IP10 can be a valuable target for MS.

NF-*κ*B is extensively present in the cytosol and the nucleus, to a lesser extent in the extracellular space, but not in the plasma membrane (Source: UniProtKB/Swiss-Prot). No small molecule or antibody is currently under clinical study for a direct blockade of NF-*κ*B, since it is inhibited by I*κ*B proteins in cytoplasm.

Clinical assays trying to inhibit NF-*κ*B have so far focused on its upstream regulators. The phosphorylation of I-*κ*B by the IKK (cat) kinase complex marks I-*κ*B for destruction via the ubiquitination pathway, thereby allowing the activation of the NF-*κ*B complex (Iwai 2012). Different research groups tried to inhibit undesired NF-*κ*B activity at several regulatory levels (Calzado et al. 2007). For example, inhibitors of IKKB-beta (or IKBKB: Inhibitor Of Nuclear Factor Kappa B Kinase Subunit Beta) aim at blocking the kinase which phosphorylates inhibitors of NF-kappa-B on two critical serine residues. Several small molecules antagonists targeting IKBKB are in phase I, II and III clinical trials for several diseases (source: Open Target (Carvalho-Silva et al. 2018b)).

Downstream of NF-*κ*B, glucocorticoids receptors (GR) also constitute an interesting research direction. Ligand-bound GR is able to antagonize the activity of immunogenic transcription factors such as nuclear factor-*κ*B (NF-*κ*B)3, AP-14,5, and T-bet6; resulting in a potent attenuation of inflammation (Hudson et al. 2018).

Altogether, these clinical assays for IP10 and NF-*κ*B pathway inhibitors strengthen the potential of the pair as MS targets, where their simultaneous inhibition lowers the risk for MS.

## 4 Discussion

We study gene-gene interactions for a number of disease maps related to multiple sclerosis. Nonetheless, the pipeline we describe here can be generalized to other diseases. It is based on epiGWAS, a SNP-level epistasis detection tool that we extend to the study of gene-level epistasis. Within each disease map, we obtained a number of significant interactions that formed novel subnetworks. Notably, we have shown complementarity between two different SNP-to-gene mappings: eQTL mapping and physical mapping. We identified 4 gene interactions mediated by potential function modifying variants. Among these interactions we retrieve one known direct binding interaction between GLI-I and SUFU, involved in oligodendrocyte precursor cells differentiation in MS. We also identified 25 gene interactions mediated by eQTLs, in particular a IP10-NFKB interaction where each gene separately has a known impact on MS. We show that the epistasis mechanism probably pass through the known regulation of IP10 transcription by NFKB. These observations validate that epistasis analysis can reveal biological interactions and confort the use of this methodology to predict new biology. To the best of our knowledge, our work is the first application of an epistasis detection tool to a specific disease which is followed by an in-depth statistical analysis and biological interpretation of the results. Nonetheless, more biological and experimental validation is needed to confirm the discovered interactions.

## 5 Data access

This study makes use of data generated by the Wellcome Trust Case-Control Consortium. A full list of the investigators who contributed to the generation of the data is available from www.wtccc.org.uk. Funding for the project was provided by the Wellcome Trust under award 076113, 085475 and 090355.

## Footnotes

I available for download from https://www.fda.gov/media/80100/download

