## Supplementary material for "A systematic analysis of gene-gene interaction in multiple sclerosis"

### A systematic analysis of gene-gene interaction in multiple sclerosis
### Supplementary Materials

#### A Distribution of SNPs in MS disease maps

Table 1: SNP and gene distributions in each disease map for eQTL and physical mappings

| internal ID | Physical mapping | | | eQTL mapping | | |
|---|---|---|---|---|---|---|
| | #SNPs | #genes | average #SNPs per gene | #SNPs | #genes | average #SNPs per gene |
| 3302 | 416 | 21 | 19.81 | 833 | 19 | 43.84 |
| 3305 | 70 | 10 | 7.00 | 238 | 8 | 29.75 |
| 3306 | 383 | 21 | 18.24 | 869 | 19 | 45.74 |
| 4455 | 755 | 38 | 19.87 | 1813 | 36 | 50.36 |
| 4593 | 1295 | 24 | 53.96 | 1647 | 17 | 96.88 |
| 4693 | 544 | 34 | 16.00 | 912 | 27 | 33.78 |
| 4703 | 331 | 28 | 11.82 | 999 | 27 | 37.00 |
| 4791 | 252 | 24 | 10.50 | 1264 | 23 | 54.96 |
| 4794 | 84 | 15 | 5.60 | 331 | 12 | 27.58 |
| 4843 | 984 | 32 | 30.75 | 1401 | 29 | 48.31 |
| 4846 | 1318 | 36 | 36.61 | 1555 | 32 | 48.59 |
| 4901 | 1173 | 35 | 33.51 | 1209 | 24 | 50.38 |
| 5199 | 656 | 28 | 23.43 | 1320 | 32 | 41.25 |
| 5288 | 515 | 27 | 19.07 | 724 | 22 | 32.91 |
| 5378 | 257 | 22 | 11.68 | 907 | 22 | 41.23 |
| 5398 | 141 | 21 | 6.71 | 1050 | 24 | 43.75 |
| 5518 | 392 | 29 | 13.52 | 1474 | 27 | 54.59 |
| 5601 | 348 | 28 | 12.43 | 742 | 25 | 29.68 |
| 5611 | 224 | 22 | 10.18 | 906 | 24 | 37.75 |

### B Visualization of epiGWAS results on MetaCore disease maps

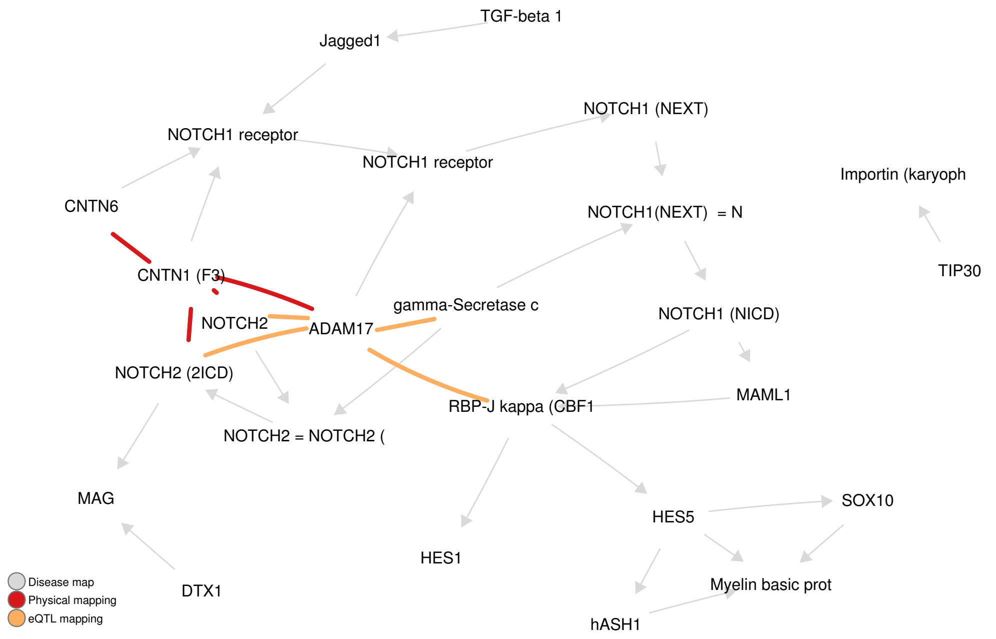


(a) DM 3302: Notch signaling in oligodendrocyte precursor cell differentiation in multiple sclerosis

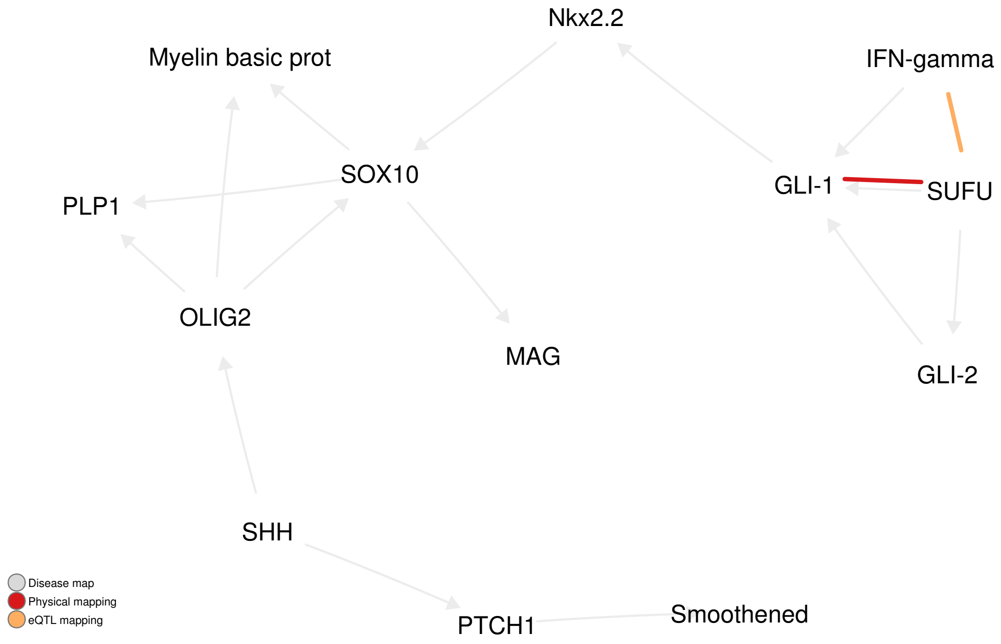


(b) DM 3305: SHH signaling in oligodendrocyte precursor cells differentiation in multiple sclerosis

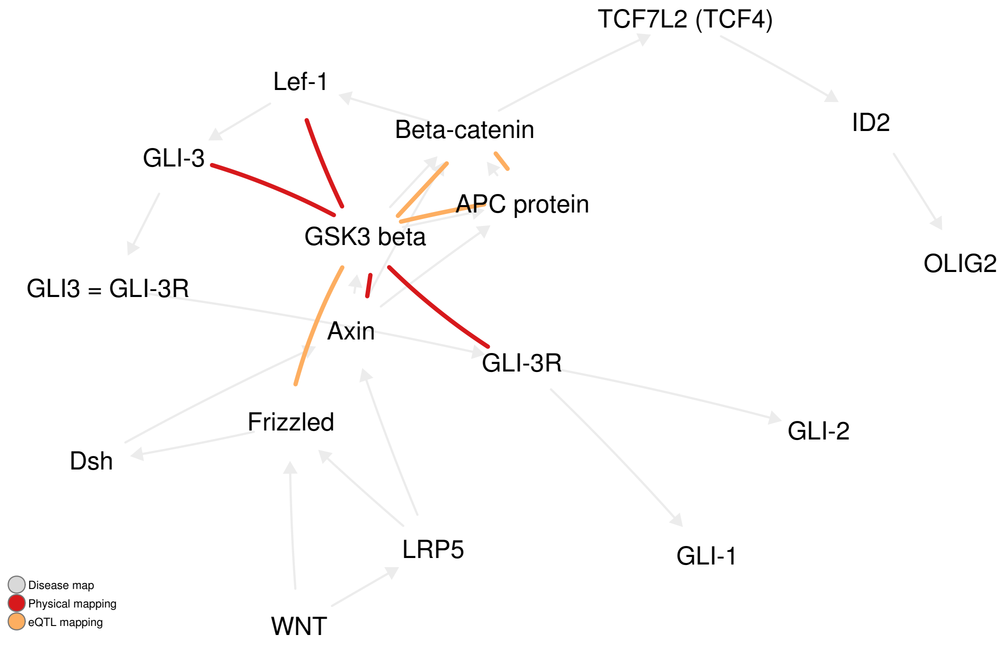


(c) DM 3306: Inhibition of oligodendrocyte precursor cells differentiation by Wnt signaling in multiple sclerosis

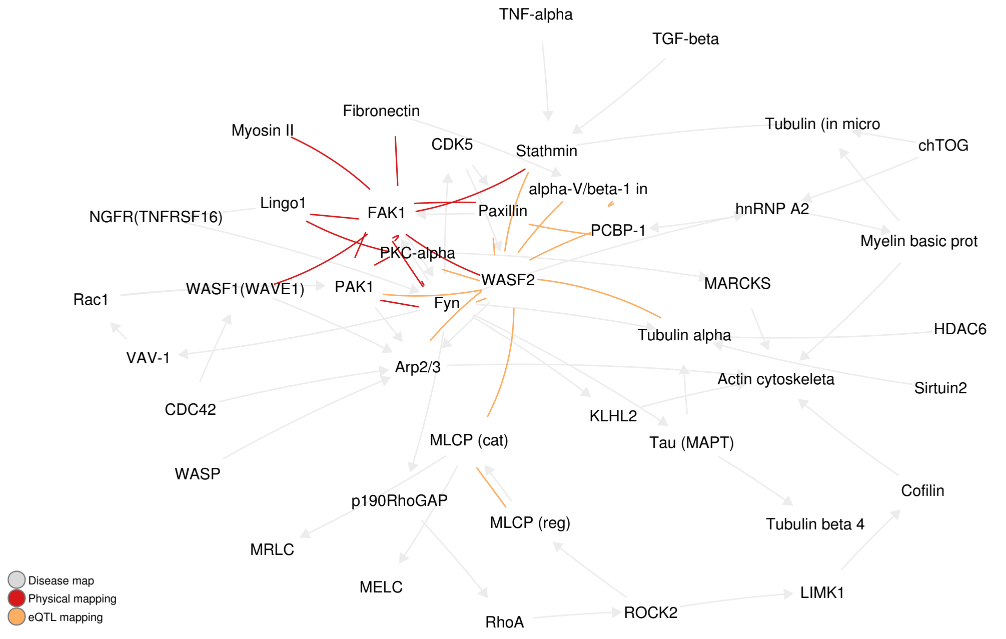


(d) DM 4455: Inhibition of remyelination in multiple sclerosis: regulation of cytoskeleton proteins

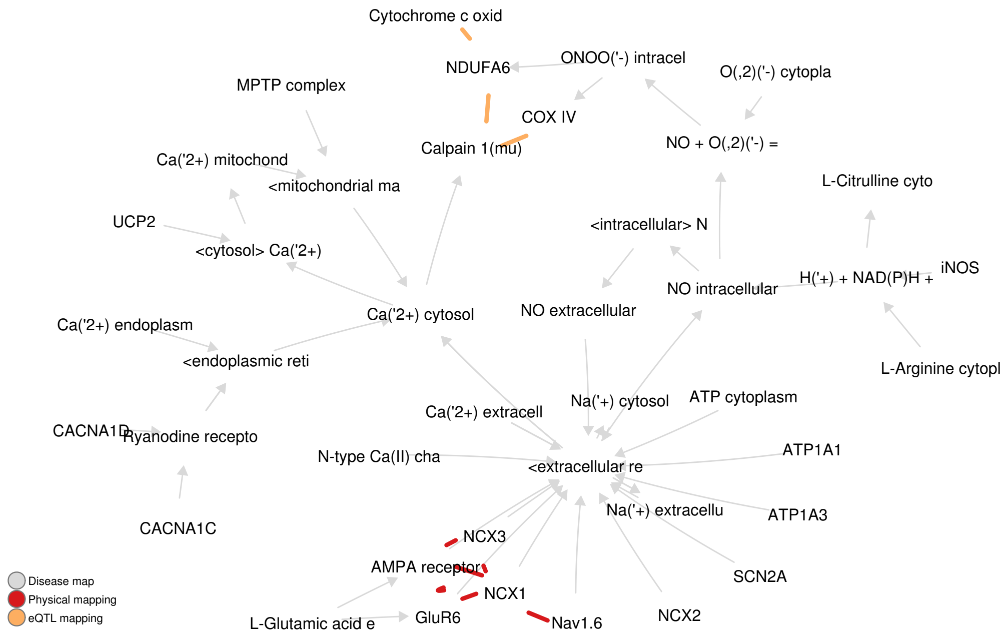


(e) DM 4593: Axonal degeneration in multiple sclerosis

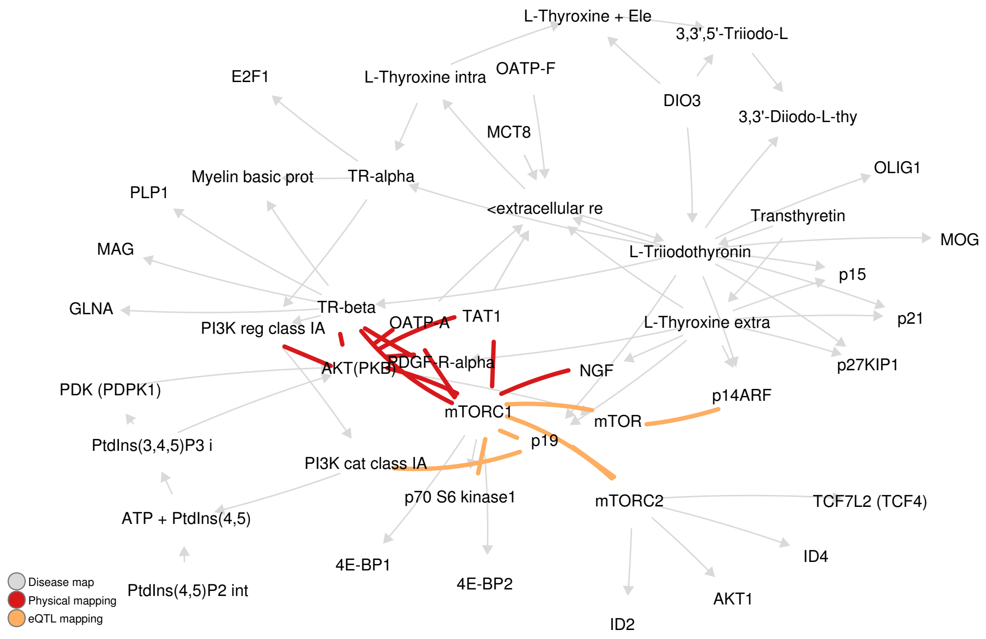


(f) DM 4693: Role of Thyroid hormone in regulation of oligodendrocyte differentiation in multiple sclerosis

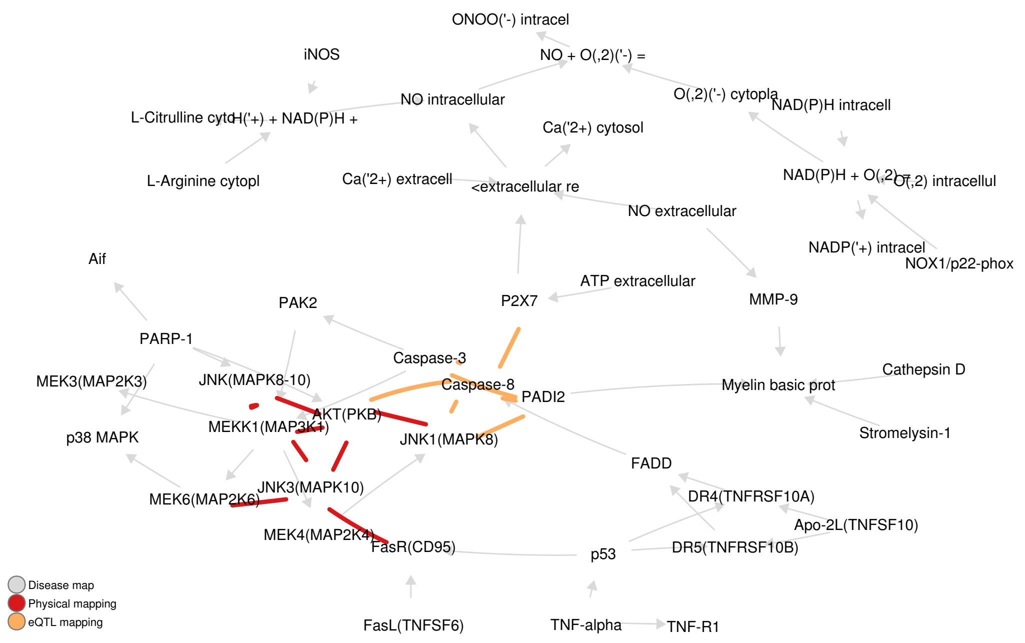


(g) DM 4703: Demyelination in multiple sclerosis

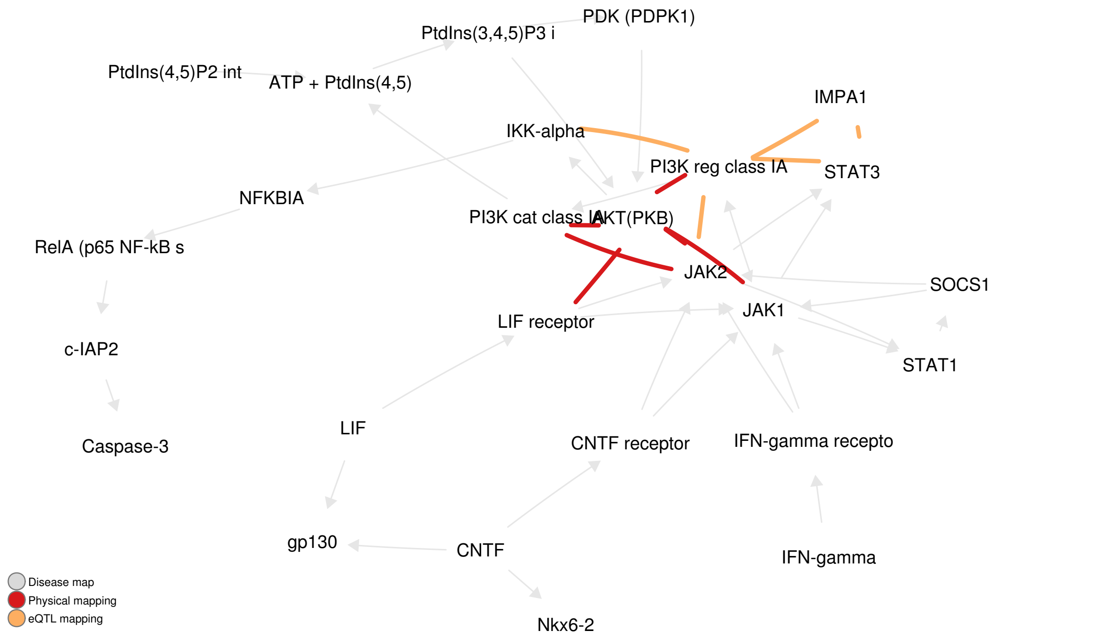


(h) DM 4791: Role of CNTF and LIF in regulation of oligodendrocyte development in multiple sclerosis

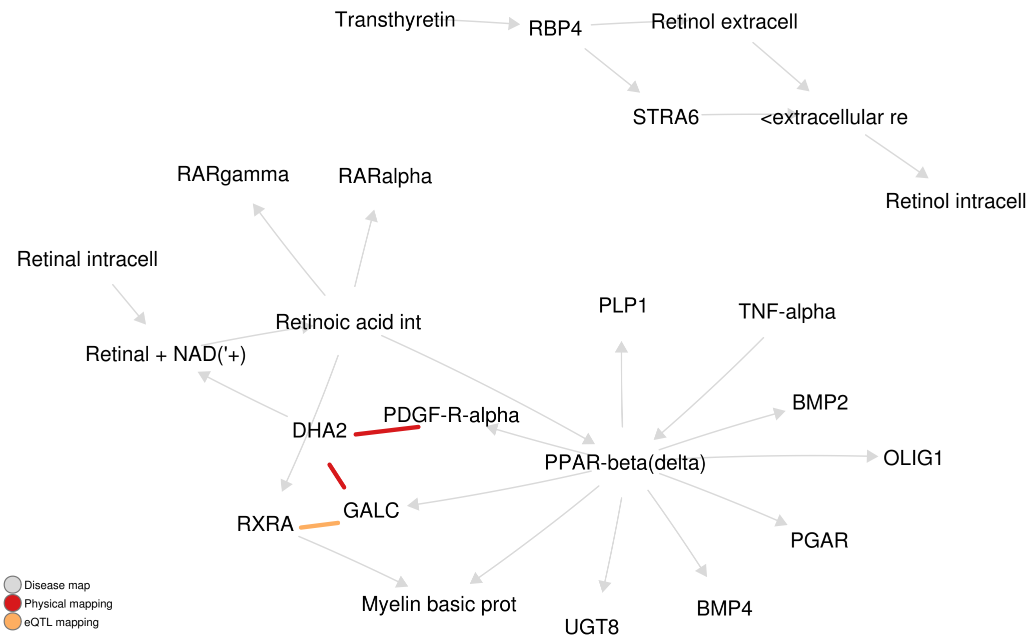


(i) DM 4794: Retinoic acid regulation of oligodendrocyte differentiation in multiple sclerosis

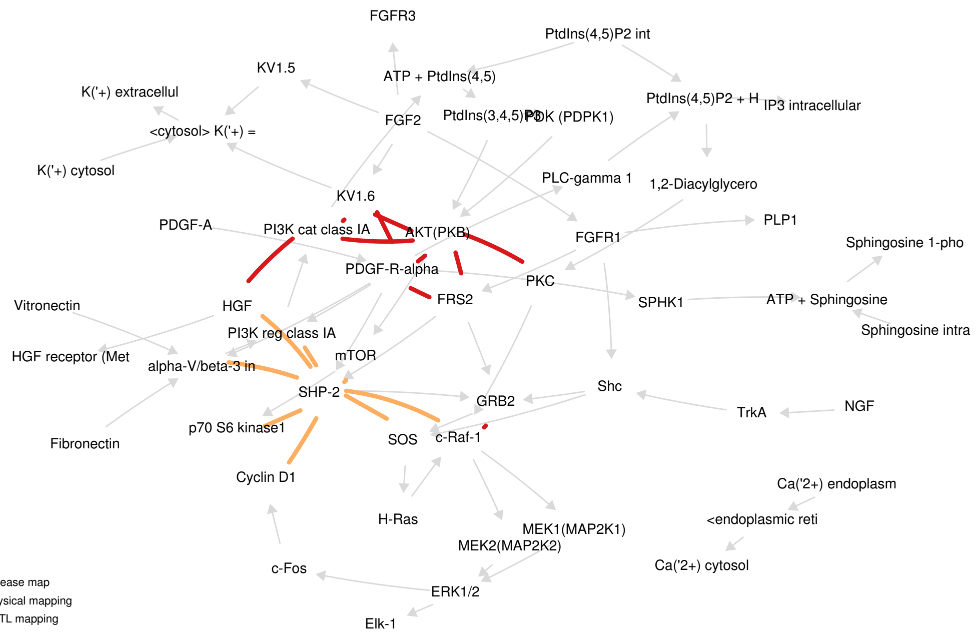


(j) DM 4843: Growth factors in regulation of oligodendrocyte precursor cells proliferation in multiple sclerosis

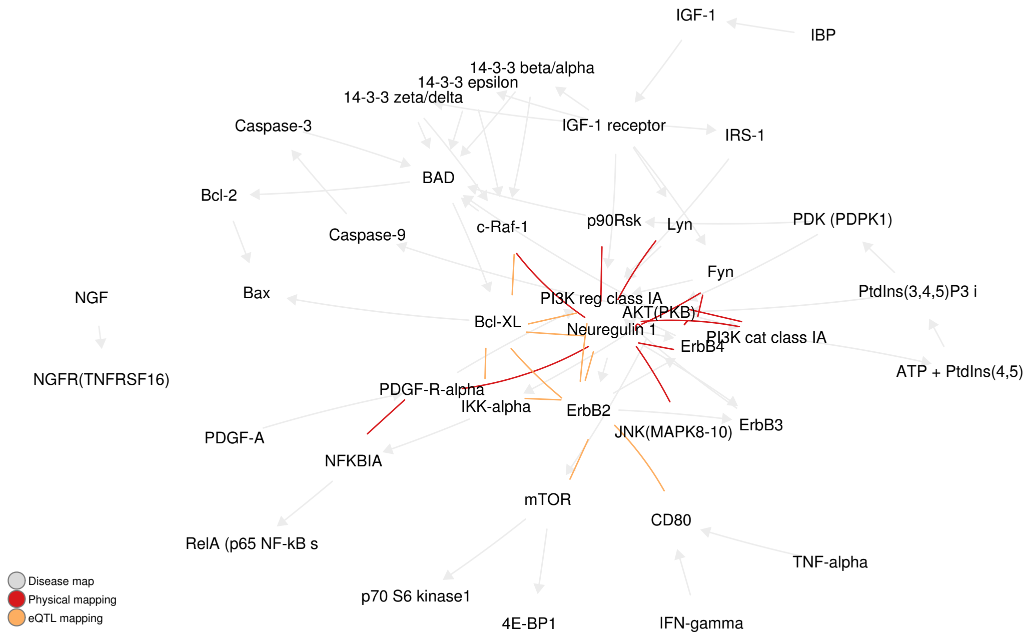


(k) DM 4846: Growth factors in regulation of oligodendrocyte precursor cells survival in multiple sclerosis

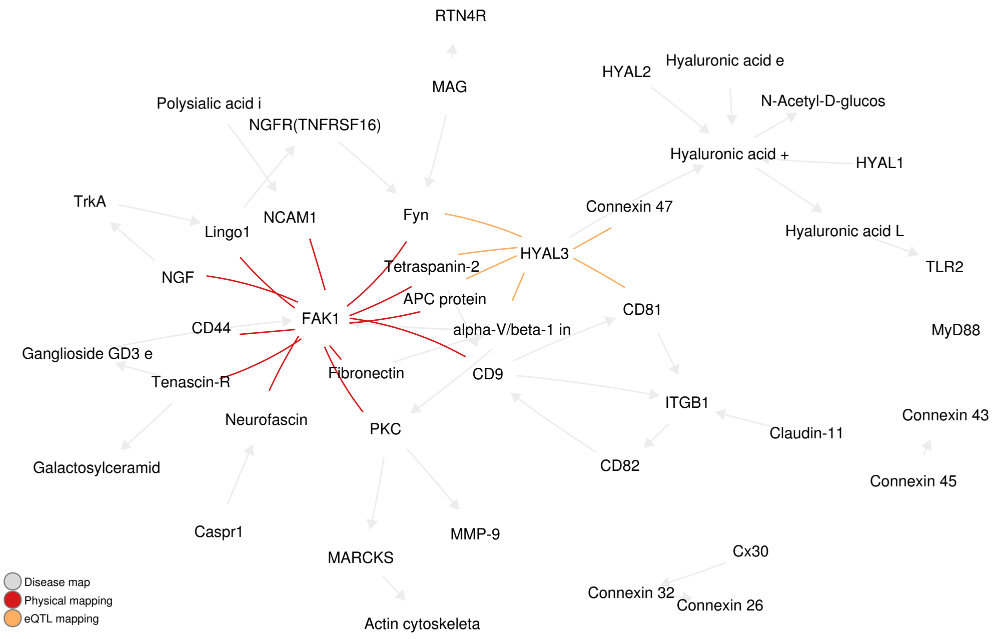


(l) DM 4901: Inhibition of remyelination in multiple sclerosis: role of cell-cell and ECM-cell interactions

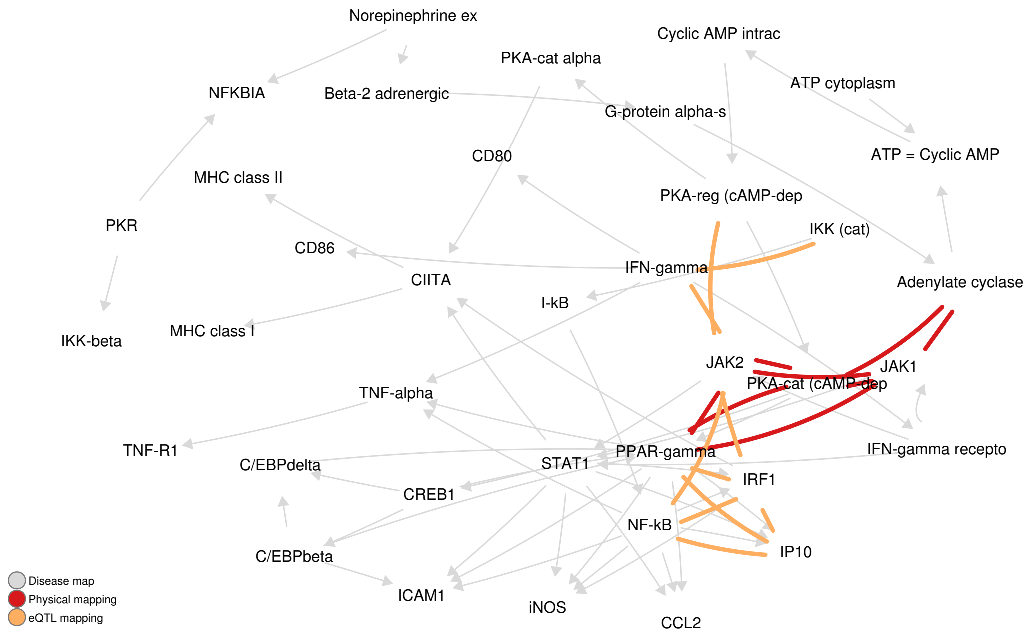


(m) DM 5199: Cooperative action of IFN-$\gamma$ and TNF-$\alpha$ on astrocytes in multiple sclerosis

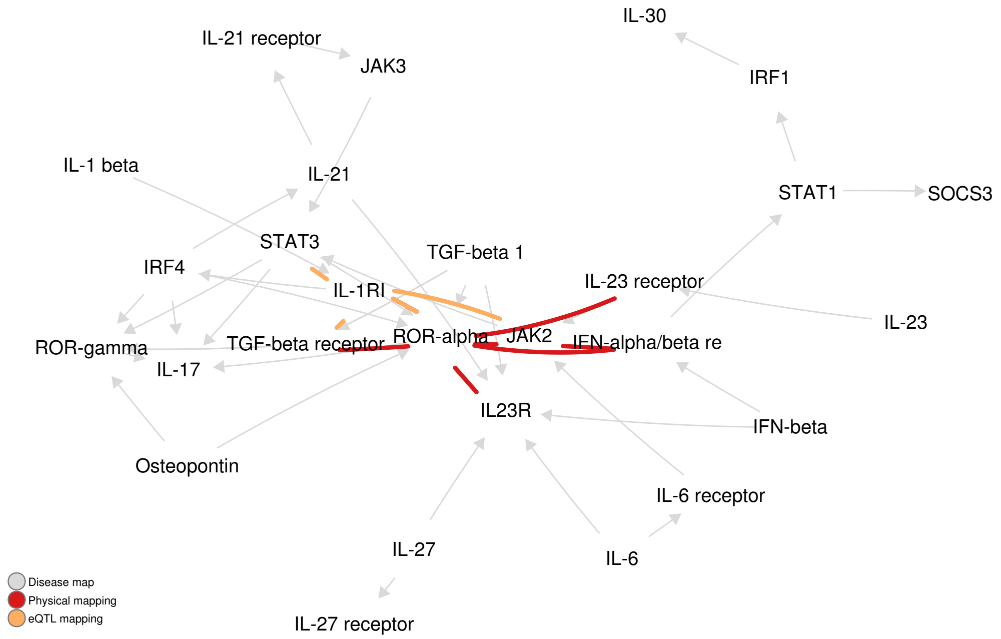


(n) DM 5288: Impaired inhibition of Th17 cell differentiation by IFN-$\beta$ in multiple sclerosis

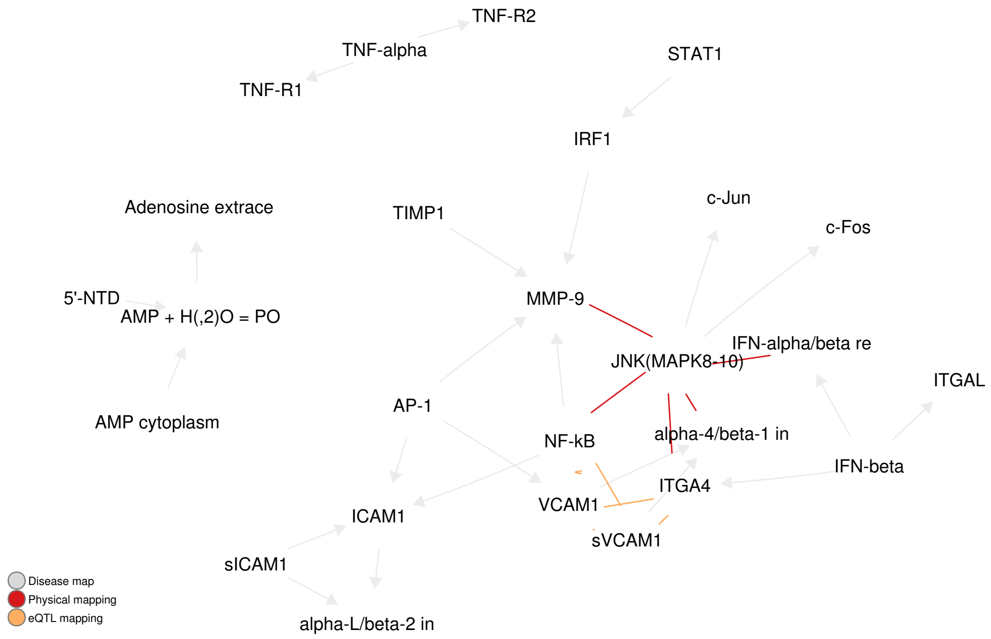


(o) DM 5378: Role of IFN-$\beta$ in the improvement of blood-brain barrier integrity in multiple sclerosis

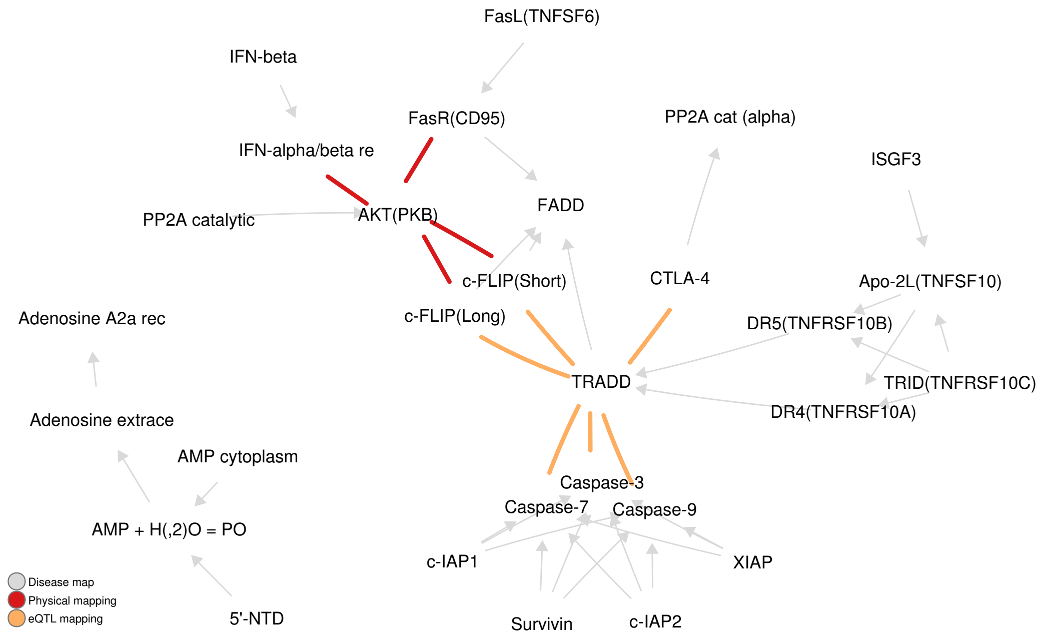


(p) DM 5398: Role of IFN-$\beta$ in activation of T cell apoptosis in multiple sclerosis

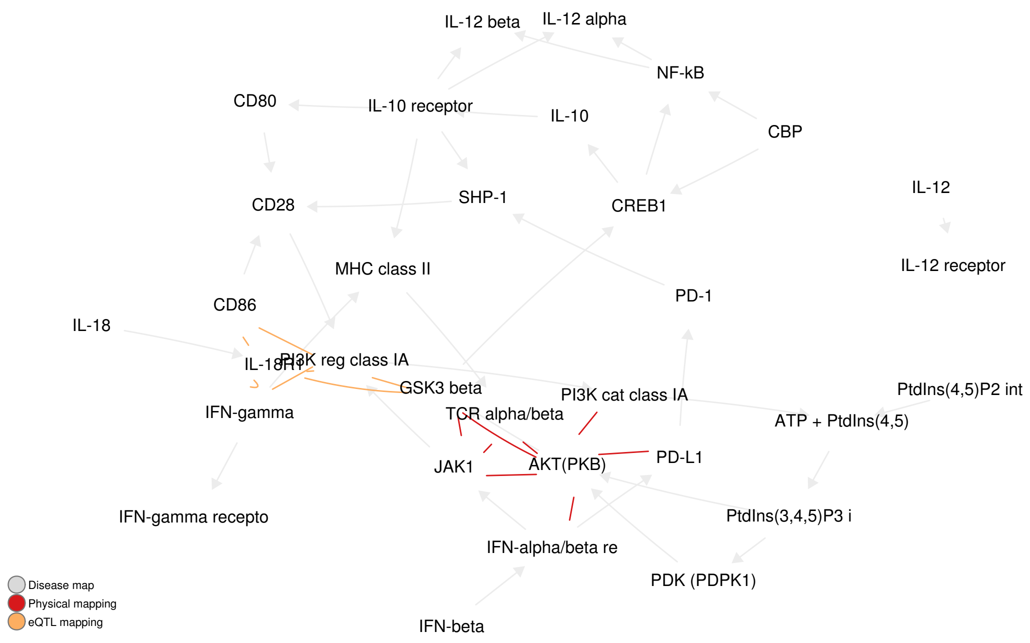


(q) DM 5518: Role of IFN-$\beta$ in inhibition of Th1 cell differentiation in multiple sclerosis

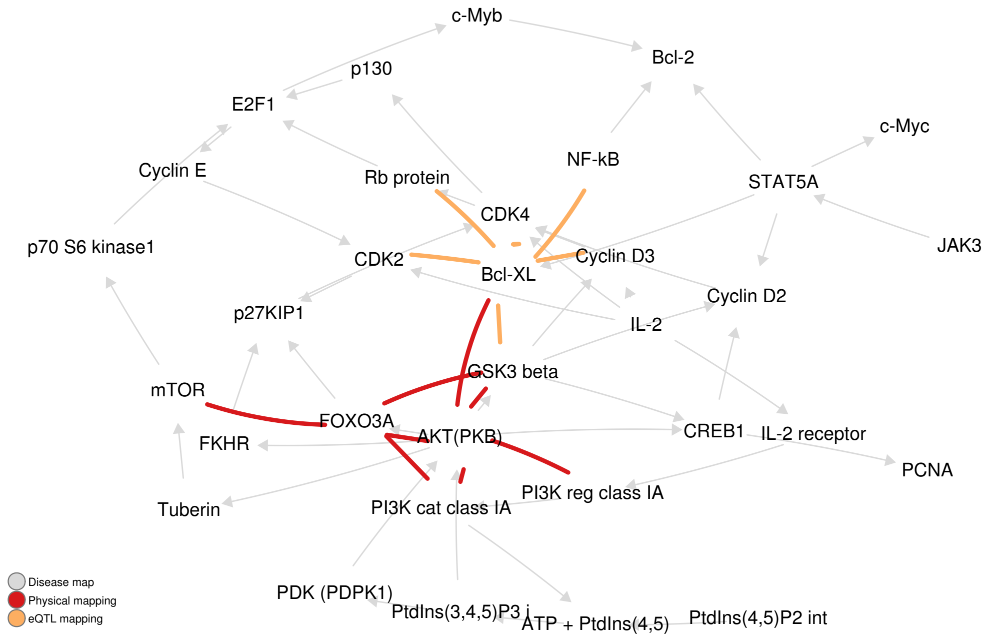


(r) DM 5601: IL-2 as a growth factor for T cells in multiple sclerosis

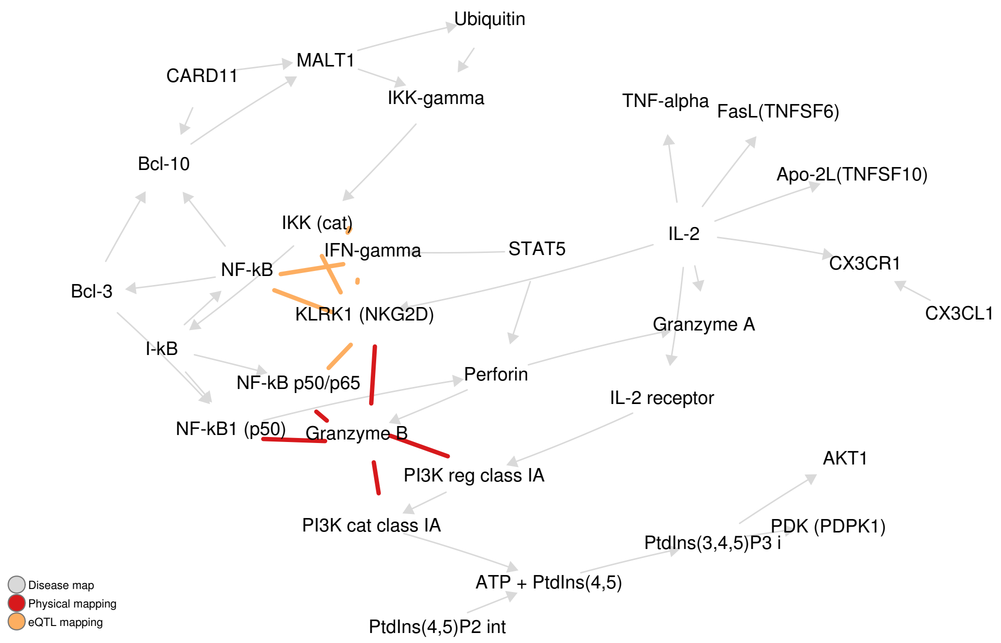


(s) DM 5611: Role of IL-2 in the enhancement of NK cell cytotoxicity in multiple sclerosis

Figure 1: Representation of the 2% top-scoring interactions for physical and eQTL mappings on the original disease maps.

### C Statistical significance of the observed network characteristics

Table 2: Enrichment analysis results for four network characteristics: connectedness, complementarity, centrality and commonality.

| internal ID | Connectedness | | Complementariy | | | Centrality | Commonality | | |
|---|---|---|---|---|---|---|---|---|---|
| | physical mapping | eQTL mapping | 1 vertex | 2 vertices | 3 vertices | maximal degree | 1 edge | 2 edges | 3 edges |
| 3302 | 0.019 | 0.021 | 0.705 | 0.254 | 0.038 | 0.147 | 0.542 | 0.149 | 0.022 |
| 3305 | 1.000 | 1.000 | 0.294 | 0.013 | 0.000 | 1.000 | 0.354 | 0.035 | 0.000 |
| 3306 | 0.061 | 0.060 | 0.792 | 0.335 | 0.054 | 0.001 | 0.761 | 0.357 | 0.099 |
| 4455 | 0.000 | 0.000 | 0.988 | 0.905 | 0.679 | 0.000 | 0.822 | 0.499 | 0.224 |
| 4593 | 0.001 | 0.014 | 0.392 | 0.056 | 0.004 | 0.724 | 0.368 | 0.072 | 0.007 |
| 4693 | 0.000 | 0.000 | 0.728 | 0.314 | 0.072 | 0.000 | 0.619 | 0.246 | 0.069 |
| 4703 | 0.000 | 0.000 | 0.649 | 0.208 | 0.033 | 0.750 | 0.479 | 0.126 | 0.020 |
| 4791 | 0.008 | 0.011 | 0.778 | 0.340 | 0.070 | 0.407 | 0.728 | 0.333 | 0.096 |
| 4794 | 0.161 | 1.000 | 0.241 | 0.011 | 0.000 | 1.000 | 0.233 | 0.028 | 0.001 |
| 4843 | 0.000 | 0.000 | 0.836 | 0.477 | 0.171 | 0.014 | 0.551 | 0.179 | 0.037 |
| 4846 | 0.000 | 0.000 | 0.938 | 0.699 | 0.361 | 0.000 | 0.782 | 0.447 | 0.191 |
| 4901 | 0.000 | 0.002 | 0.947 | 0.729 | 0.391 | 0.000 | 0.561 | 0.187 | 0.040 |
| 5199 | 0.000 | 0.000 | 0.726 | 0.291 | 0.057 | 0.134 | 0.785 | 0.439 | 0.183 |
| 5288 | 0.004 | 0.014 | 0.791 | 0.366 | 0.082 | 0.009 | 0.697 | 0.307 | 0.090 |
| 5378 | 0.012 | 0.011 | 0.673 | 0.215 | 0.026 | 0.341 | 0.561 | 0.172 | 0.030 |
| 5398 | 0.012 | 0.004 | 0.779 | 0.346 | 0.074 | 0.052 | 0.567 | 0.178 | 0.034 |
| 5518 | 0.002 | 0.002 | 0.723 | 0.283 | 0.050 | 0.251 | 0.665 | 0.275 | 0.074 |
| 5601 | 0.001 | 0.002 | 0.847 | 0.472 | 0.146 | 0.032 | 0.713 | 0.338 | 0.109 |
| 5611 | 0.004 | 0.004 | 0.687 | 0.245 | 0.037 | 0.405 | 0.602 | 0.210 | 0.048 |

### D Content of resulting subnetworks in therapeutic targets

Table 3: Number of drug targets in the resulting subnetworks for each disease map and its statistical significance.

| internal ID | Number of included drug targets | $p$-value |
|---|---|---|
| 3302 | 0 | 1.000 |
| 3305 | 0 | 1.000 |
| 3306 | 1 | 0.378 |
| 4455 | 2 | 0.380 |
| 4593 | 6 | 0.009 |
| 4693 | 2 | 0.382 |
| 4703 | 0 | 1.000 |
| 4791 | 2 | 0.154 |
| 4794 | 1 | 0.222 |
| 4843 | 2 | 0.808 |
| 4846 | 2 | 0.500 |
| 4901 | 2 | 0.265 |
| 5199 | 1 | 0.875 |
| 5288 | 2 | 0.728 |
| 5378 | 4 | 0.024 |
| 5398 | 2 | 0.347 |
| 5518 | 4 | 0.275 |
| 5601 | 1 | 0.768 |
| 5611 | 0 | 1.000 |

### E MetaCore disease maps

#### E.1 Disease map 3305

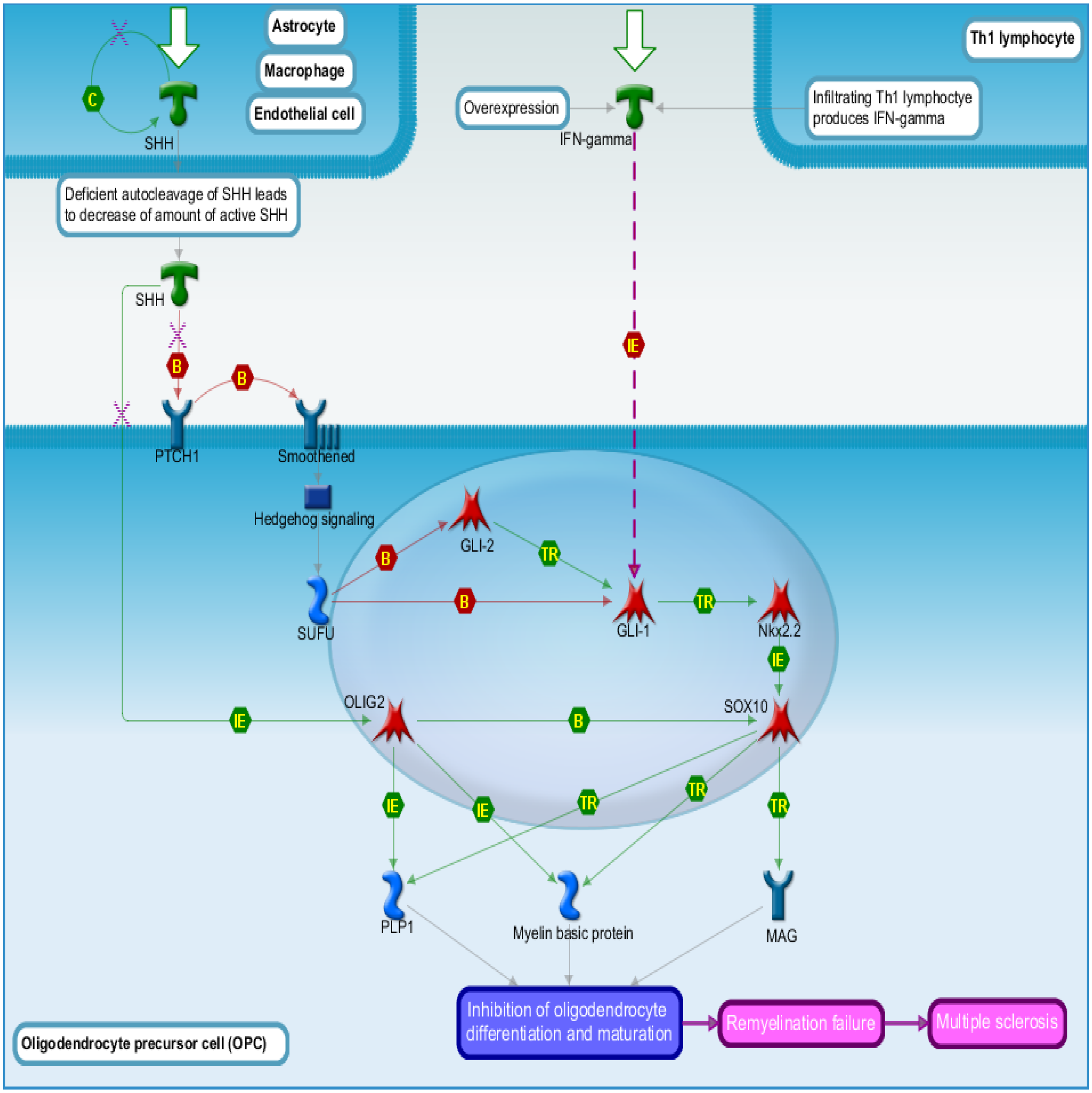


Figure 2: Sonic Hedgehog signaling in oligodendrocyte precursor cells differentiation in multiple sclerosis (DM 3305).

### E.2 Disease map 4455

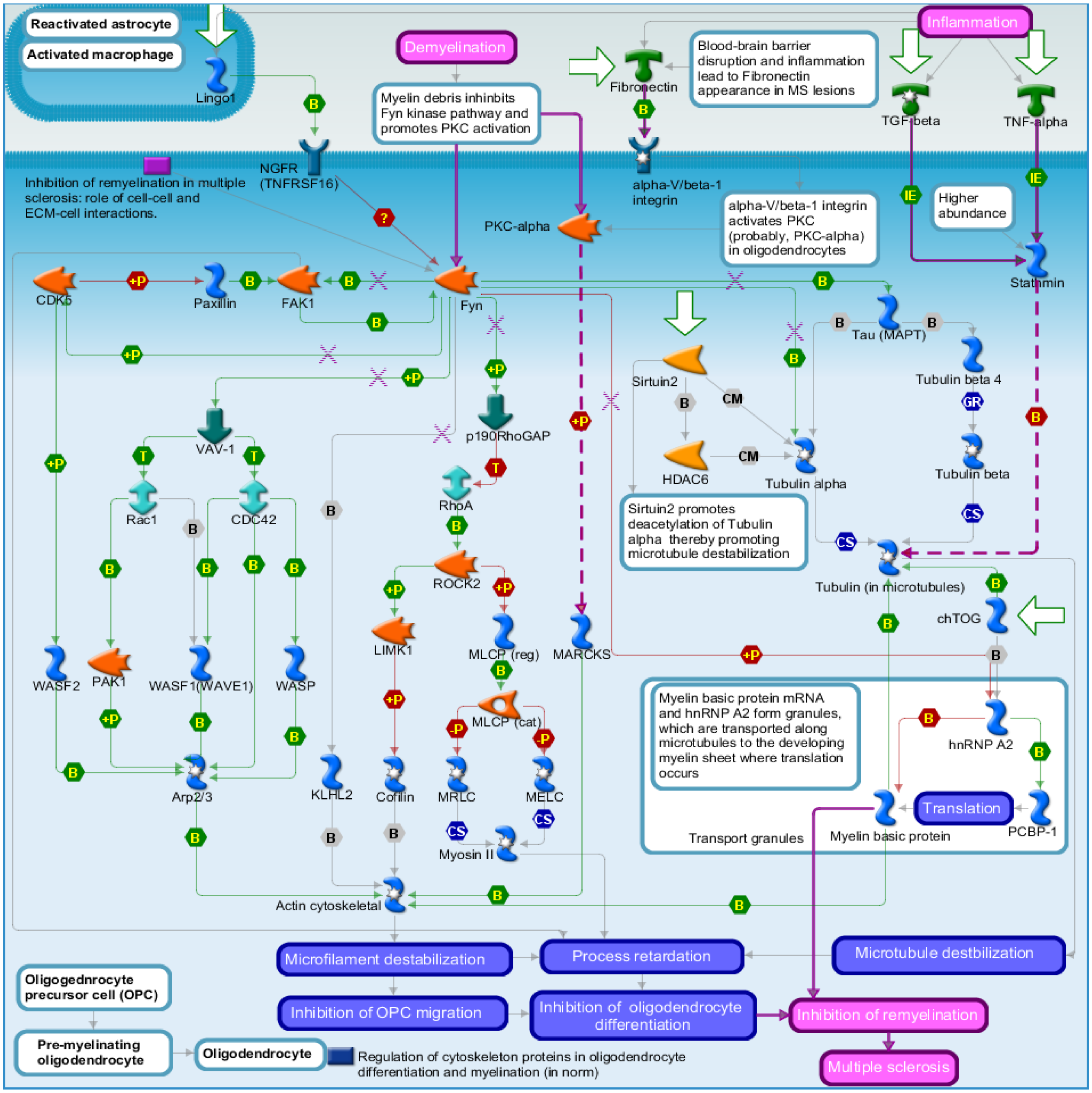


Figure 3: Inhibition of remyelination in multiple sclerosis: regulation of cytoskeleton proteins (DM 4455).

#### E.3 Disease map 5199

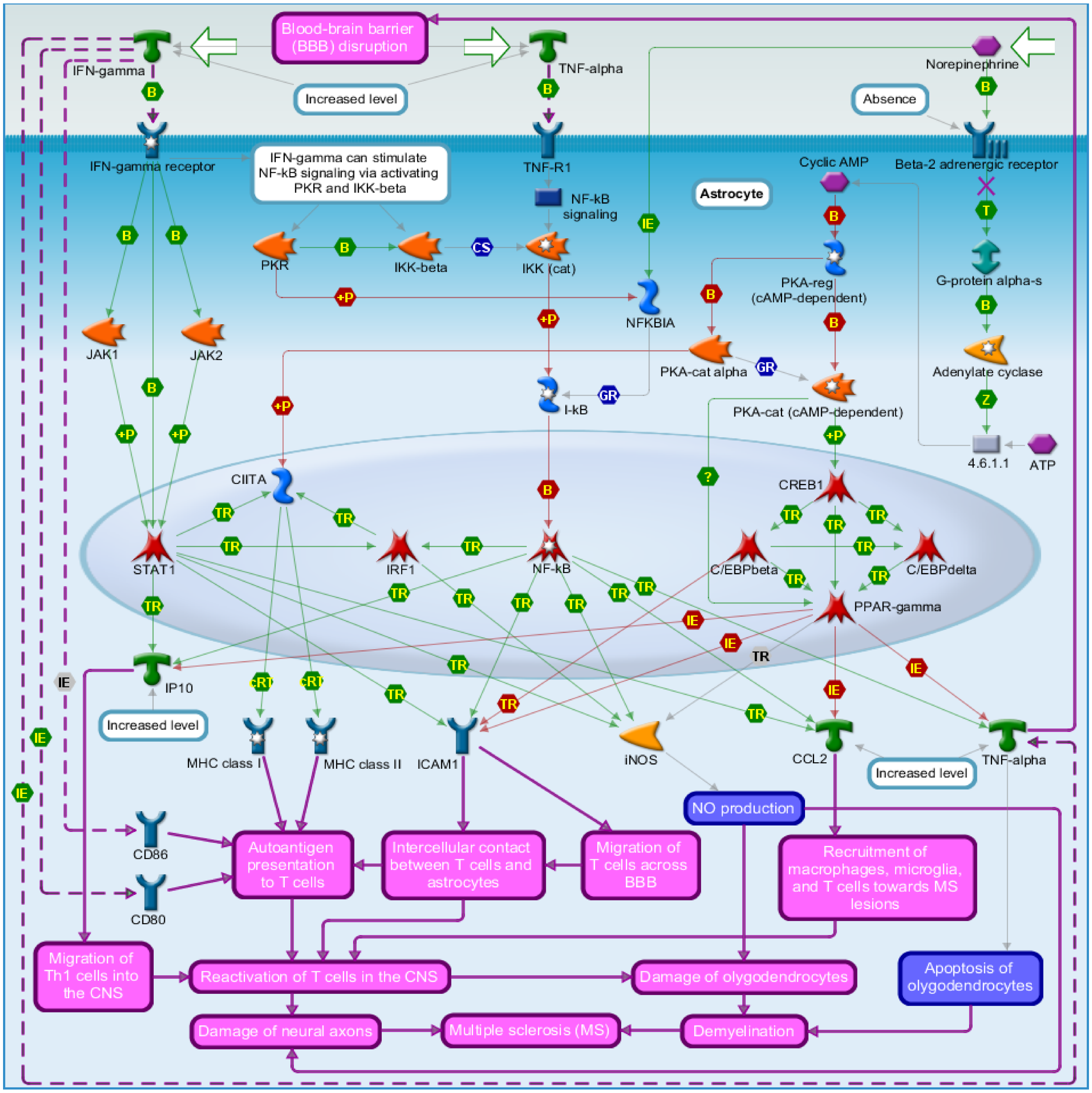


Figure 4: Cooperative action of IFN-gamma and TNF-alpha on astrocytes in multiple sclerosis (DM 5199).

### F Filtering pipeline

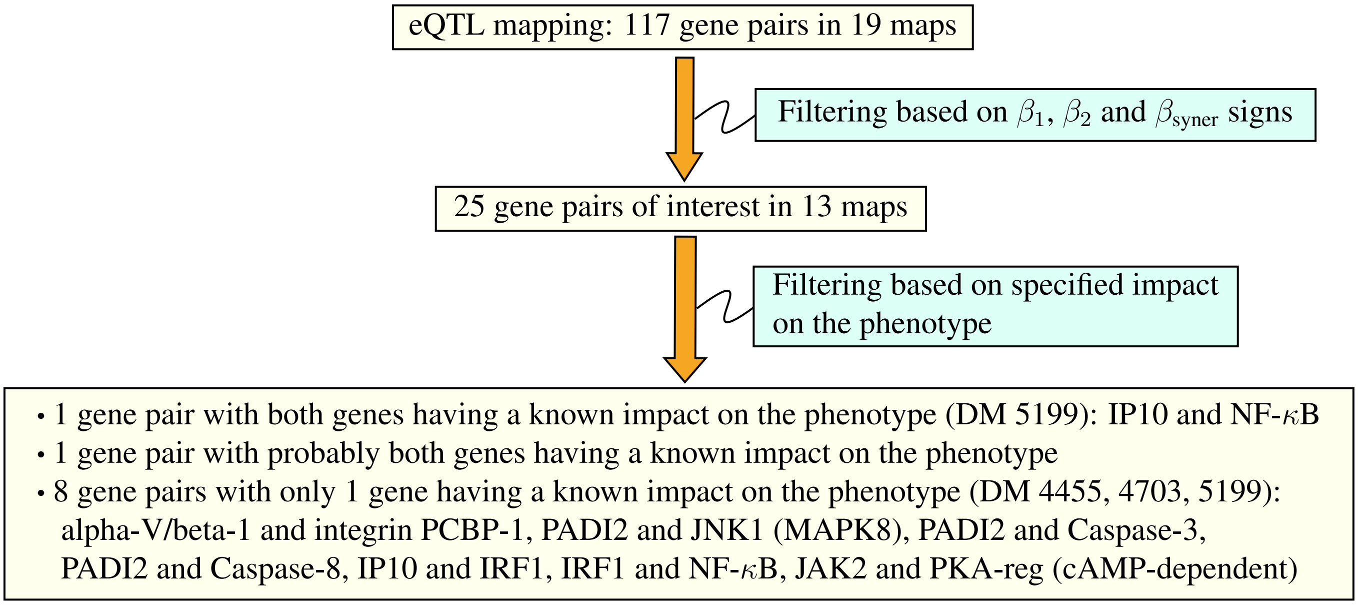


Figure 5: Filtering process for gene pairs identified by eQTL mapping.

### G Physical mapping

Table 4: Pairs of genes identified by physical mapping, and selected on the basis of their SNPs' consequence as a protein dysfunction.

| internal ID | Gene pair | Type of interaction |
|---|---|---|
| 3305 | GLI-1 and SUFU | direct interaction between the genes, but unspecified impact on MS |
| 4703 | AKT (PKB) and MEKK1 (MAP3K1) | no direct interaction between the genes, but AKT has a specified impact on MS |
| 5611 | Granzy me B and KLRK1 (NKG2D) | no direct interaction between the genes, and unspecified impact on MS |
| | Granzyme B and PI3K cat class IA | no direct interaction between the genes, and unspecified impact on MS |

### H eQTL mapping

Table 5: Compiled results of gene pairs identified by epistasis, and filtered according to the scheme in Fig 2, with their specified or unknown impact on MS.

| internal ID | Title | Interacting gene pair | | $\beta_x$ | $\beta_y$ | $\beta_{\text{syner}}$ | Specified impact on MS (activation or inhibition) |
|---|---|---|---|---|---|---|---|
| 3302 | Notch signaling in oligodendrocyte precursor cell differentiation in multiple sclerosis | RBP-J kappa (CBF1) | ADAM17 | 1.40 | 1.37 | 0.02 | no |
| 3305 | SHH signaling in oligodendrocyte precursor cells differentiation in multiple sclerosis | | | | | | |
| 3306 | Inhibition of oligodendrocyte precursor cells differentiation by Wnt signaling in multiple sclerosis | Beta-catenin | GSK3 beta | 1.27 | 1.84 | 0.00 | no |
| 4455 | Inhibition of remyelination in multiple sclerosis: regulation of cytoskeleton proteins | alpha-V/beta-1 integrin | PCBP-1 | 1.27 | 0.96 | 0.01 | probably yes for alpha-V/beta-1 integrin |
| 4593 | Axonal degeneration in multiple sclerosis | | | | | | |
| 4693 | Role of Thyroid hormone in regulation of oligodendrocyte differentiation in multiple sclerosis | | | | | | |
| 4703 | Demyelination in multiple sclerosis | PADI2 | JNK1(MAPK8) | -1.42 | -1.58 | -0.02 | PADI2 enhances in disease |
| 4703 | Demyelination in multiple sclerosis | PADI2 | Caspase-3 | -1.56 | -1.96 | -0.01 | PADI2 enhances in disease |
| 4703 | Demyelination in multiple sclerosis | PADI2 | Caspase-8 | -1.47 | -1.21 | -0.03 | PADI2 enhances in disease |
| 4703 | Demyelination in multiple sclerosis | JNK1(MAPK8) | Caspase-8 | -1.58 | -1.21 | -0.01 | no |
| 4791 | Role of CNTF and LIF in regulation of oligodendrocyte development in multiple sclerosis | IMPA1 | STAT3 | 1.41 | 1.10 | 0.02 | no |
| 4791 | Role of CNTF and LIF in regulation of oligodendrocyte development in multiple sclerosis | PI3K reg class IA | STAT3 | 1.40 | 1.10 | 0.05 | no |

| | | | | | | | |
|---|---|---|---|---|---|---|---|
| 4794 | Retinoic acid regulation of oligodendrocyte differentiation in multiple sclerosis | | | | | | |
| 4843 | Growth factors in regulation of oligodendrocyte precursor cells proliferation in multiple sclerosis | alpha-V/beta-3 integrin | SHP-2 | 1.34 | 1.97 | 0.07 | no |
| 4843 | Growth factors in regulation of oligodendrocyte precursor cells proliferation in multiple sclerosis | SHP-2 | c-Raf-1 | 1.63 | 1.63 | 0.09 | no |
| 4846 | Growth factors in regulation of oligodendrocyte precursor cells survival in multiple sclerosis | ErbB2 | Neuregulin 1 | 1.10 | 1.58 | 0.11 | no |
| 4846 | Growth factors in regulation of oligodendrocyte precursor cells survival in multiple sclerosis | Neuregulin 1 | Bcl-XL | -1.49 | -1.17 | -0.02 | no |
| 4901 | Inhibition of remyelination in multiple sclerosis: role of cell-cell and ECM-cell interactions | Fyn | HYAL3 | -1.99 | -1.38 | -0.07 | no |
| 5199 | Cooperative action of IFN-$\gamma$ and TNF-$\alpha$ on astrocytes in multiple sclerosis | IP10 | IRF1 | 1.41 | 1.12 | 0.09 | yes for IP10 |
| 5199 | Cooperative action of IFN-$\gamma$ and TNF-$\alpha$ on astrocytes in multiple sclerosis | IP10 | NF-$\kappa$B | 1.39 | 0.98 | 0.09 | yes for both genes |
| 5199 | Cooperative action of IFN-$\gamma$ and TNF-$\alpha$ on astrocytes in multiple sclerosis | IRF1 | NF-$\kappa$B | 1.16 | 0.88 | 0.07 | yes for NF-$\kappa$B |
| 5199 | Cooperative action of IFN-$\gamma$ and TNF-$\alpha$ on astrocytes in multiple sclerosis | JAK2 | PKA-reg (cAMP-dependent) | 1.14 | 1.25 | 0.02 | yes for JAK2 |
| 5288 | Impaired inhibition of Th17 cell differentiation by IFN-beta in multiple sclerosis | IL-1RI | ROR-alpha | -1.16 | -1.29 | -0.09 | yes (probable) |
| 5378 | Role of IFN-beta in the improvement of blood-brain barrier integrity in multiple sclerosis | | | | | | |
| 5398 | Role of IFN-beta in activation of T cell apoptosis in multiple sclerosis | CTLA-4 | TRADD | -1.61 | -2.61 | -0.04 | no |
| 5398 | Role of IFN-beta in activation of T cell apoptosis in multiple sclerosis | Caspase-3 | TRADD | -1.96 | -2.21 | -0.07 | no |

| | | | | | | | |
|---|---|---|---|---|---|---|---|
| 5518 | Role of IFN-beta in inhibition of Th1 cell differentiation in multiple sclerosis | IFN-$\gamma$ | PI3K reg class IA | 1.29 | 1.40 | 0.07 | no |
| 5518 | Role of IFN-beta in inhibition of Th1 cell differentiation in multiple sclerosis | GSK3 beta | IL-18R1 | 1.39 | 1.36 | 0.02 | no |
| 5518 | Role of IFN-beta in inhibition of Th1 cell differentiation in multiple sclerosis | PI3K reg class IA | CD86 | -0.96 | -1.13 | -0.18 | no |
| 5601 | IL-2 as a growth factor for T cells in multiple sclerosis | GSK3 beta | Bcl-XL | -0.85 | -1.17 | -0.03 | no |
| 5611 | Role of IL-2 in the enhancement of NK cell cytotoxicity in multiple sclerosis | | | | | | |
